# Degenerate DropSynth for Simultaneous Assembly of Diverse Gene Libraries and Local Designed Mutants

**DOI:** 10.1101/2023.12.11.569291

**Authors:** Andrew S. Holston, Samuel R. Hinton, Kyra A. Lindley, Nora C. Kearns, Calin Plesa

## Abstract

Protein engineering efforts often involve the creation of hybrid or chimeric proteins, where functionality critically hinges on the precise design of linkers and fusion points. Traditional methods have been constrained by a focus on single genes or the random selection of fusion points. Here we introduce an approach which enables the creation of large gene libraries where each library comprises a multitude of diverse, specifically designed genes, each with a corresponding set of programmatically designed fusion points or linkers. When combined with multiplex functional assays, these libraries facilitate the derivation of generalized engineering principles applicable across whole protein families or domain types. Degenerate DropSynth is a multiplex gene synthesis technique which allows for the assembly of up to eight distinct variants for each of the 1,536 designed parent genes in a single reaction. We assemble chimeric sensor histidine kinases and demonstrate the assembly of genes up to 1 kbp in length with an 8% rate of perfect assemblies per gene. Our findings indicate that incorporating an increased number of variants in droplets containing barcoded beads does not significantly affect the rate of perfect assemblies. However, maintaining a consistent level of degeneracy across the library is important to ensure good coverage and reduce inequality. The results suggest the potential for scaling this process to assemble at least 8,000 distinct variants in a single reaction. Degenerate DropSynth enables the systematic exploration of protein families through large-scale, programmable assembly of chimeric proteins, moving beyond the limitations of individual protein studies.

## Introduction

Protein engineering seeks to design proteins with novel properties and functions by determining the sequences corresponding to a particular targeted function or property. Its potential applications are immense and span from drug development and diagnostics to biofuel production and environmental remediation. One important aspect of protein engineering is the creation of chimeric or hybrid proteins^1–4^. These proteins, especially prevalent in cellular signaling^5^, are constructed by combining different modules to create new or enhanced functionalities. This is particularly useful in the development of biomass saccharification^6^, cellular signaling^5^, and complex biosynthesis pathways^7^ to name a few. Central to the success of these chimeric proteins are the design and engineering of fusion points and linkers. The choice of fusion point can significantly impact the function of the resulting chimeric protein, as it can influence the spatial arrangement of the protein domains and their ability to interact with each other and with other molecules^1,8^. Linkers, on the other hand, are connectors that provide flexibility and space between the domains, enabling proper folding and independent functioning^9,10^. Their characteristics (length, composition, and conformation) significantly affect the activity, stability, and solubility of the chimeric protein^9,11^. Thus, the strategic selection of fusion points and linkers is a key element in the field of protein engineering.

Although most protein engineering efforts in this area have focused on single chimeric proteins, an increasingly desirable approach would be to decipher the rules for engineering entire protein families rather than individual proteins. Towards this end a potential Design-Build-Test-Learn strategy could consist of: (1) designing large amounts of diverse relevant hybrids through metagenomic mining or rational computational approaches, (2) assembling large libraries of specifically designed variants spanning many diverse genes, (3) functionally characterizing the library using a multiplexed functional assay (4) feeding the resulting data into computational or machine learning (ML) models which can discern the underlying patterns affecting functionality, (5) repeating this process using computational or ML generated variants and feeding the results back into the model until some target threshold for accuracy is achieved. This manuscript tackles the second step in this process and demonstrates how to build large libraries of designed local mutant variants for many diverse genes.

Previous methods to generate fusion or linker variants are unable to generate designed variants for a diverse library of gene fusions. Methods utilizing nuclease mediated truncation such as SHIPREC^12^ and ITCHY^13^ create many fusions at random points. Although PCR primer based methods such as PATCHY^14^ allow control over the fusion points, they are only applicable to single genes and do not work with diverse gene libraries, due to the lack of conserved sequence in the priming region among different library members.

We previously introduced a multiplex gene synthesis method, DropSynth 2.0, and demonstrated that it was capable of assembling 1,536 genes with a median 501 bp in length in a single reaction with 64% coverage and a 25% percent perfects at the amino acid level^15^ (∼15% at nucleotide level). Although this approach increased the assembly scale significantly, in the context of protein engineering, given the immense number of variants possible, creating a large number of variants for many diverse genes would quickly saturate the 1536x genes possible in a typical reaction. Scaling would require the use of many separate libraries. As such, we sought to see if multiple variants for the same parent sequence could be synthesized within each droplet. In this approach, DropSynth proceeds exactly as before, Fig. 1a, with oligos processed to expose a 12-nt single stranded barcoded overhang, which is then hybridized and ligated on to corresponding barcoded beads, followed by emulsification into droplets, and emulsion polymerase cycling assembly (ePCA). The degenerate DropSynth approach leverages the fact that any additional oligo targeted to a barcode which contains the same ePCA overhangs as the gene will also participate in the assembly reaction (Fig. 1b) and create a variant of the gene in a programmable manner. This approach provides several advantages. First, the marginal cost of each additional variant is only the cost of an additional oligo as all other reagents remain unchanged. Second, this provides a simple path towards much higher scales without larger sets of barcoded beads. Although in this study we focus on variants at the end of each gene, such that only the last oligo is modified, we note that this approach could easily be modified to create internal defined variants (Sup. Fig S1), combinatorial variants (Sup. Fig S2), or even multiple different full-length genes if completely orthogonal overlaps are used. While background-isolation from complex pools is critical to prevent cross-hybridization and proper assembly, small numbers of genes can be successfully assembled together with minimal screening for othogonality^16^.

**Fig. 1.**
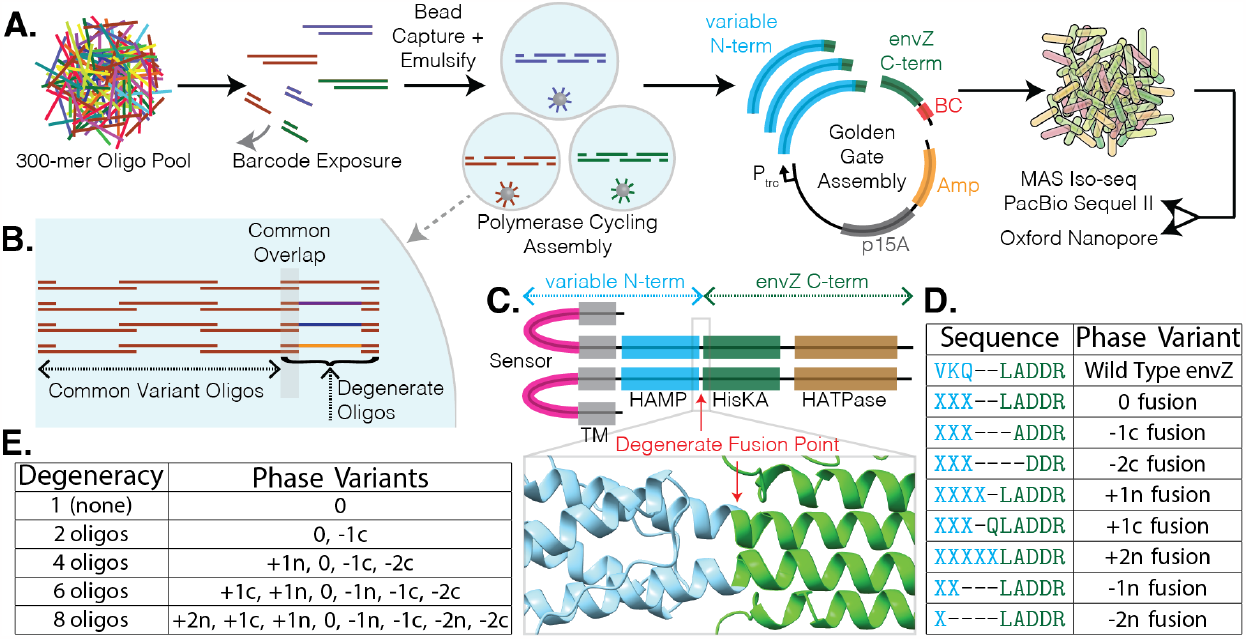
Degenerate DropSynth for rational fusion point engineering: **a**. Overview of the DropSynth protocol and cloning scheme. **b**. Within each barcoded microbead droplet, multiple versions of the last oligo are added. These degenerate oligos share a common overlap and can participate in the assembly reaction, with each encoding a different mutant variant. **c**. Domain architecture of a typical sensor histidine kinase. We assembled SHKs with diverse sensory and HAMP domains each linked to the C-terminal portion of the well characterized histidine kinase envZ. Fusion points were generated below the HAMP domain. (bottom insert) AlphaFold2 predicted structure around the fusion site of a NarX-EnvZ chimera. Proper function of these chimeric sensors requires the correct phase orientation between the fused alpha helices. **d**. By adding or removing residues from the N-terminal or C-terminal fragments, the relative phase orientation between the alpha helices on either side of the fusion point is changed. **e**. We tested assembly reactions where genes contained between 1 (no degeneracy) and 8 degenerate oligos encoding different phase variants to assess its impact on the gene assembly process.

We apply the degenerate DropSynth approach to the creation of chimeric sensor histidine kinases (SHK) as a proof of concept. SHKs are one of the most abundant protein families found in nature with millions of receptors that can sense a wide variety of stimuli including small molecules^17,18^, light^19^, pH^20^, metal ions, and osmotic pressure^21^. These homodimeric modular proteins typically contain an extracellular sensing domain, transmembrane domains, and signaling domains which help propagate the activation signal to a kinase domain^22^ (Fig. 1c). Phosphorylation of a histidine residue is transferred to the aspartate residue on a specific response regulator protein which can then dimerize and activate transcription of a target promoter^23^. Despite their abundance, activating ligands are only known for a small number of SHKs^18^ since individual characterization is a slow and laborious process. Large scale characterization and deorphanization of this family could be achieved if the modular sensory domains could be swapped onto a well characterized kinase domain to allow for multiplexed testing of many receptors^24,25^. Many challenges exist in such an approach, as detailed elsewhere^26–29^. One issue is the selection of a fusion point due to the diversity and uncertainty in the exact domain boundaries as detected by HMMs or other methods, which makes it difficult to create functional chimeras, even if the signal transduction mechanism of both halves is the same. This makes it difficult to put the upstream (helix) portion into the proper phase orientation with the downstream (helix) portion^30–34^. In this work we focus on building chimeric fusions in a region just below the HAMP signaling domain, a homodimeric four alpha-helix parallel coiled-coil region. We use degenerate DropSynth to create many phase variants for each sensor domain through the controlled addition or subtraction of amino acid residues on either the C or N terminal fragments of each chimera as shown in Fig. 1d. Any variant can be made as it is programmatically encoded on a corresponding assembly oligo, with the only requirements being that the variant sequence length can fit onto the corresponding oligo and be placed outside the common overlap region.

As a proof of concept, in each library, we tested the assembly of between 1 and 8 different phase variants for each gene, as shown in Fig. 1e. In other words, in the same reaction, some droplets would only assemble one gene with no additional variants (degeneracy of 1), while at the other extreme droplets would assemble 8 different variants of the same gene. This allowed us to control for inter-reaction variability from factors such as the DNA recovery from the emulsions, PCR amplification, and processing yields. We designed two sets of proteins based on their length. One with 1,530 proteins to be assembled with 4x 300-mer oligos and another with 1,531 proteins to be assembled with 5x 300-mer oligos. We made two different codon libraries for each set for a total of four libraries with 6,122 total genes distributed among them. We have previously shown that using multiple codon versions increases the chances of a successful assembly. With the extra degenerate variants added, the four libraries encoded for a total of 10,862 proteins and 21,724 genes with distribution of degeneracy levels shown in Sup. Fig. S3. In the two 4-oligo libraries, we distributed the microbead barcodes relatively uniformly among the different degeneracy levels (median 292 microbead barcodes, s.d. 22), while in the 5-oligo libraries over half of the microbead barcodes had no degeneracy (level 1) (Sup. Fig. S4), to ensure sufficient statistics for the longer length. The use of 300-mer oligos allowed us to reach assembly lengths of 1 kbp with 5 oligos, effectively doubling the length demonstrated previously with DropSynth 2.0^15^.

## Results and Discussion

Increased levels of degenerate oligos have a minimal impact on the percentage perfects of assembled genes. The four libraries were successfully assembled (Fig. 2ab) using the standard DropSynth protocol, Golden Gate cloned into an randomly barcoded expression vector^35^, and long-read sequenced using both MAS ISO-seq^36^ (PacBio Sequel II) and Oxford Nanopore. Among genes with at least 100 barcodes, we found a median of 16.6% and 17.3% DNA perfects for the 4 oligo library codon 1 and codon 2 respectively (Fig. 2c-top). We explored the impact of adding increasing numbers of degenerate oligos on the rates of perfects observed. Comparing the rates of genes from the no degeneracy set to those with 2, 4, 6, or 8 we found no statistically significant differences except for Codon 2 degeneracy level 6, which showed a weakly significant decrease (p=0.014). These results suggest that the presence of additional assembly oligos has no impact on the percentage of perfects. For the five oligo assemblies we see a median of 10.2% and 13.0% DNA perfects codon 1 and codon 2 respectively (Fig. 2c-bottom). For the codon 1 library we see significant decreases for degeneracy levels of 4 (8.2%, p=0.02), 6 (7.6%, p=0.02), and 8 (6.4%, p=9E-6) relative to the no degeneracy case. A similar pattern is observed in codon 2 library with significant decreases for degeneracy levels of 4 (11.2%, p=0.0004), 6 (11.6%, p=0.0004), and 8 (11.6%, p=0.03). These decreases are attributed to the imbalance in the final representation between the variants from different degeneracy levels which is exponentially increased by PCR amplification, as discussed later, and can be observed when plotting the perfects rate against the number of barcodes observed (Sup. Fig. S5). Comparing the rates seen at the DNA level to those at the amino acid level, where synonymous mutations have been collapsed onto the parent sequence, we see a consistent 4.0% (s.d. 0.2%) higher rate at the protein level for 4 oligo assemblies and a 2.7% (s.d. 0.8%) difference for 5 oligo assemblies, as shown in Sup. Fig. S6. As expected, this suggests that the fraction of synonymous mutations decreases as length increases.

**Fig. 2.**
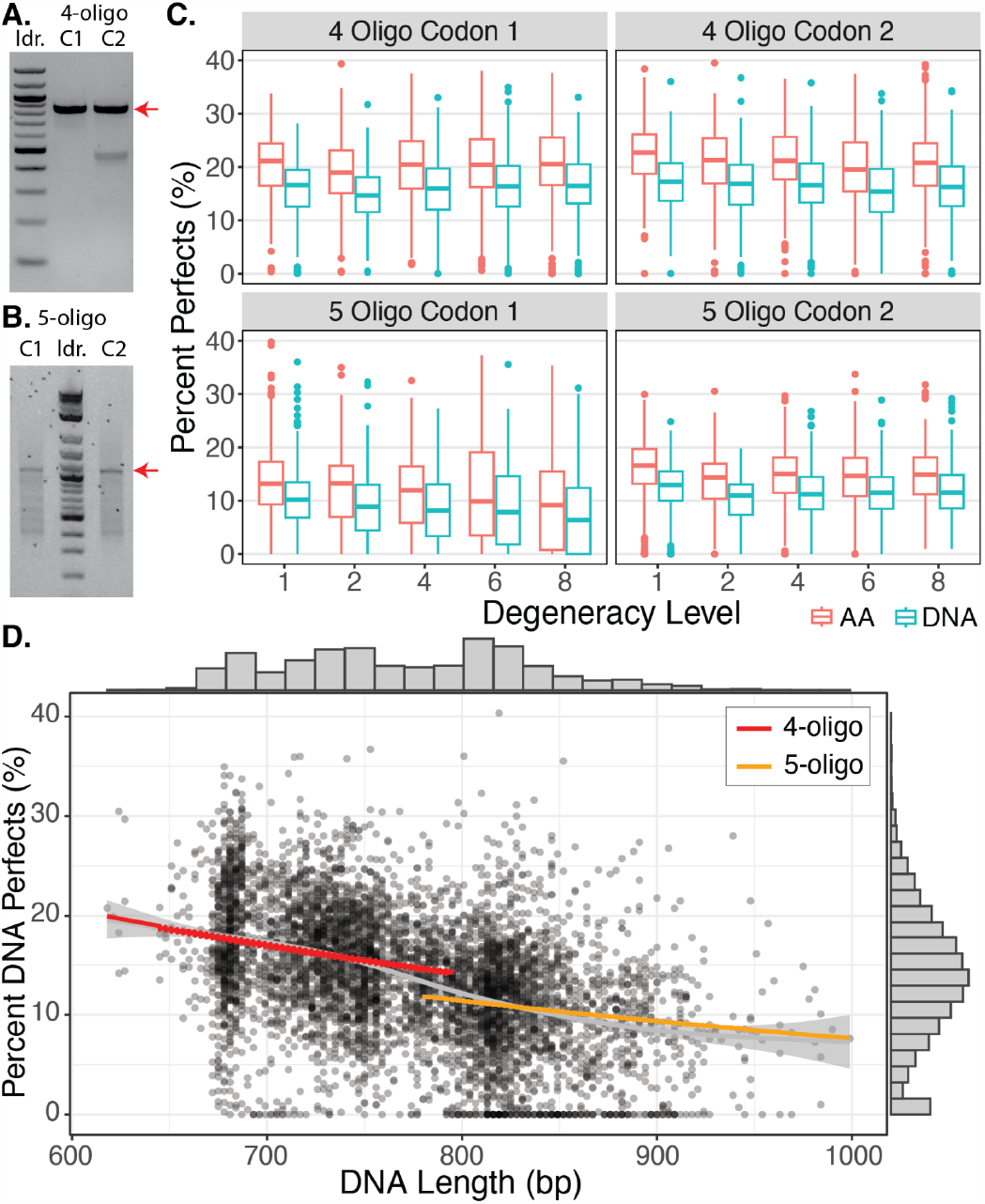
4x and 5x 300-mer oligo assemblies. **a.-b**. Assembly of the two 4-oligo and 5-oligo libraries respectively. **c**. The percentage perfects observed for variants with at least 100 barcodes shown both at the AA level (collapsed on synonymous mutations) and DNA level. **d**. The percentage of DNA perfects as a function of length showing a consistent reduction from ∼20% at 600 bp down to ∼8% by 1 kbp. Grey line is a smoothed GAM fit while the colored lines are fits for 4 and 5 oligos with the model described in the text.

Deletions are the dominant errors found in assemblies. To quantify the relative frequencies of various error types per kilobase pair (kbp), we analyzed CIGAR alignment strings. Our analysis revealed comparable frequencies for insertions (median 1.47, s.d. 0.22) and single base deletions (median 1.44, s.d. 0.25), as shown in Sup. Fig. S7. Interestingly, mismatch frequencies were similar in the 4-oligo libraries (median 1.47) but significantly higher in the 5-oligo libraries (median 4.56). Across all libraries, the occurrence of multi-base deletions was notably high (median 5.75, s.d. 2.08). Delving deeper into deletion lengths, we observed that single base deletions were most common (Sup. Fig. S8a). However, the presence of a considerable number of longer deletions indicates a higher probability that any given deletion is part of a longer multi-base deletion rather than an isolated single-base deletion (Sup. Fig. S8b). The higher rates of deletions are consistent with previous reports using microarray derived oligos^37^. Further investigation is required to determine whether these long multi-base deletions primarily originate from oligo synthesis or the ePCA process. Additionally, the elevated mismatch rates in the 5-oligo libraries could potentially be attributed to the ePCA process itself, given that other parameters like the number of cycles are comparable to those in the 4-oligo libraries, and mismatch rates should not be highly dependent on sequencing depth. We note that these numbers were derived with minimap2^38^, a general purpose aligner, as opposed to an exhaustive global alignment like Needleman-Wunsch^39^.

The rate of perfect assemblies correlates inversely with gene length, a trend we attribute to the propagation and combination of errors throughout the oligo synthesis, PCR amplification, and emulsion PCR (ePCA) assembly processes. This relationship is illustrated in Fig. 2d, where we plot the percentage of perfect assemblies for genes across all degeneracy levels as a function of their length. The 17 genes shorter than 650 bp show a median perfect assembly rate of 18.2% (s.d. 4.7%) while the 22 genes above 950 bp exhibit a rate of 8.2% (s.d. 5.2%). To understand this trend, we developed a simple model incorporating the error rates from oligo synthesis, PCR amplification, and ePCA assembly. With an error rate of 5.52E-6 errors per base per cycle^40^, Kapa HiFi amplification of the oligos and assembled genes has a relatively modest impact on the percentage of perfects. For example, Kapa HiFi based PCR amplification, performed over 27-28 cycles for oligos and 29 cycles for assembled genes, would theoretically allow 68% of 1 kbp genes to remain error-free post-PCR. For oligo synthesis with a 1 in 3000 bp error rate (based on vendor provided rates), we expect a 61% probability that all 5 oligos (300-mers) are perfect, which would reduce to 47% if the error rate increases to 1 in 2000 bp. This calculation, admittedly conservative, does not account for the selective pressure against propagation of errors in critical regions such as primers, restriction sites, and overlaps during the assembly process. By combining these factors, our model suggests an overall estimated perfect assembly rate of 41% for 1 kbp genes. Yet, the actual observed rate of perfect assemblies at this length is only around 8.2%, implying that the ePCA process might be the predominant source of errors, or alternatively, the actual error rates for PCR and oligo synthesis are higher than those used in our calculations.

Decreases in observed coverage are attributed to reduced representation at higher degeneracy levels. We determined the coverage for all libraries and degeneracy levels, where coverage is defined by the number of variants for which at least one perfect protein sequence is observed. We combined both PacBio and Nanopore data to maximize the sequencing depth of our data. For the 4 oligo libraries we observe 84% and 71% coverage for a degeneracy level of 1, which reduces to 63% and 45% by degeneracy of 8, as shown in Fig. 3a. When both codon versions are combined over proteins this improves to 93% for degeneracy of 1 and 76% by degeneracy level 8. Despite an average 0.69-fold decrease in percentage coverage, this reduction is modest compared to the 8-fold increase in scale brought about by higher degeneracy. In absolute numbers, the total count of observed genes rose from 244 and 207 at degeneracy level of 1 to 1448 and 1026 by degeneracy level of 8, due to the increased scale for the 4 oligo libraries which have a roughly uniform distribution of microbead barcodes (median 292, s.d. 22) among different degeneracy levels (Sup. Fig. S4). For the 5 oligo libraries we observe 49% and 70% coverage for a degeneracy of 1, dropping to 20% and 50% by degeneracy of 8. However, combining over the two codon versions again raised coverage levels to 82% for a degeneracy of 1 and 58% for a degeneracy level of 8.

**Fig. 3.**
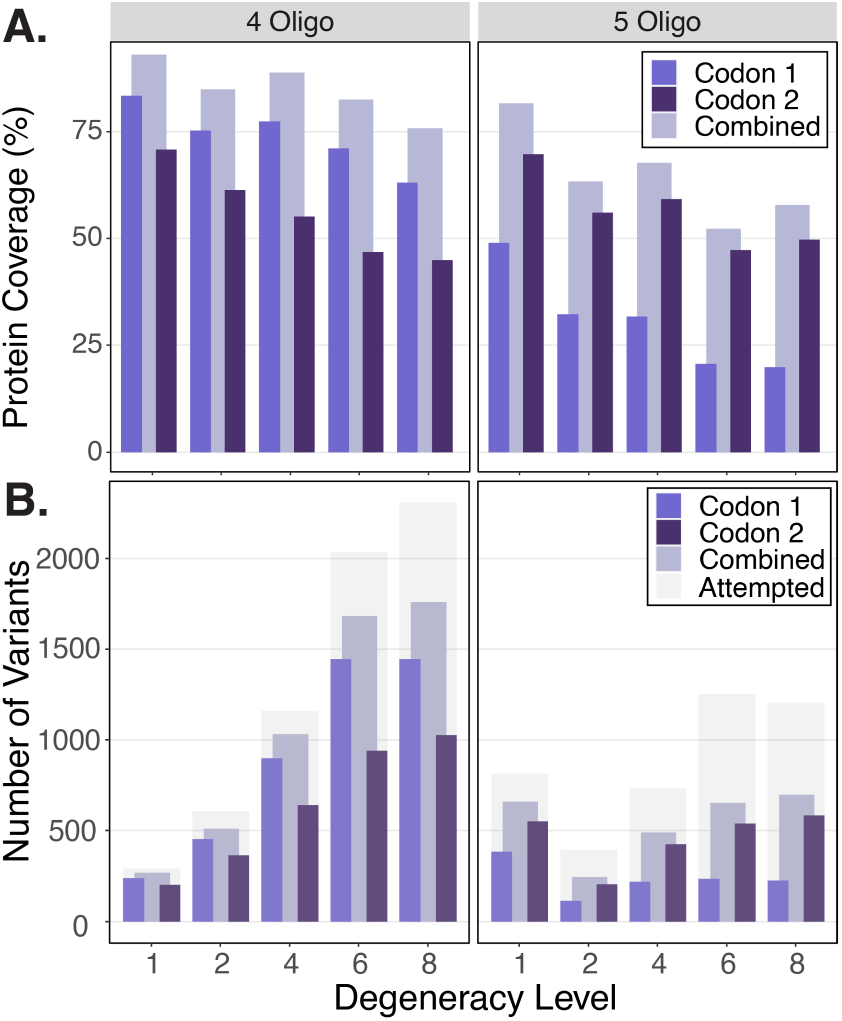
Protein coverage. The number of designed protein variants for which we see at least one perfect amino acid sequence. Coverage data are presented for each codon library, both individually and combined over the same proteins. **a**. The percentage observed relative to the number of variants designed. **b**. The absolute numbers of variants observed, contrasted against the total designed variants, which are depicted in light gray. Notably, the percent coverage decreases less than the increase in degeneracy scale, resulting in a net increase in the total number of successfully assembled protein variants.

We looked for trends which could explain this drop in coverage. In examining the fraction of barcodes observed versus the corresponding fraction of overall designs as a function of degeneracy level (Sup. Figure S9), we observe an exponential decrease with increasing degeneracy. We still see this decay when these numbers are scaled by the degeneracy level to account for lower amounts of DNA (Sup. Figure S10). We hypothesize that this trend results from the suppression PCR amplification post-assembly. Each barcoded bead in the DropSynth assembly has a limited DNA loading capacity, leading to a situation where the amount of DNA for each variant is inversely proportional to the degeneracy level. Consequently, a variant in a droplet with a degeneracy level of 8 will, on average, contain 8 times less assembled DNA than a variant with a degeneracy level of 1. This disparity is further exacerbated during suppression PCR, which significantly amplifies these initial differences. A simple PCR amplification model fits the log transformed relationship between barcodes observed per variant versus expected variant concentration relationship quite well, with R^2^ values ranging from 0.945 to 0.998 (Sup. Figure S11). Here, we calculate the expected variant concentration based on the proportion of barcoded microbeads with a given degeneracy level to the total number of variants at that level. This finding underscores the importance of maintaining a consistent degeneracy level across all variants in a degenerate DropSynth reaction.

Looking ahead, if a full 1536x library is created with a uniform degeneracy level of 8, encoding 12,288 variants, we conservatively estimate that protein coverage would be at least ∼8,000 for a single library and about 9,000 when combined over two different codon versions for 4 oligo assemblies, and roughly 6,000 for a single library and about 7,000 when combined over two codon versions for 5 oligos. These estimates are based on sufficient sequencing depth and are likely conservative, considering the potential improvements achievable with a consistent degeneracy across all library barcodes.

We determined the uniformity in the representation of the variants. The Gini coefficient, which quantifies distribution equality on a scale from 0 (perfect equality) to 1 (perfect inequality), was calculated for each degeneracy level in all four libraries (Sup. Fig. S12). This ranges from 0.71 to 0.88 (median 0.81, s.d. 0.06) for the 4 oligo libraries with a slightly increasing trend as degeneracy increases. In contrast, the 5 oligo libraries exhibited higher Gini coefficients, ranging from 0.83 to 0.93 (median 0.87, s.d. 0.04), indicating less uniformity in variant representation. These values are in line with previous observations (ranging 0.69 to 0.94), suggesting the increased degeneracy has a minor effect relative to other factors such as PCR bias.

Barcode mapping of constructs over 500 bp requires long read sequencing, beyond what can be achieved with Illumina sequencing. We investigated the use of Oxford Nanopore sequencing as an alternative to PacBio MAS-Iso-seq. Despite Oxford Nanopore’s inherently higher error rate in raw reads, previous studies have demonstrated that this can be significantly mitigated by collapsing reads onto barcodes^41,42^. To determine the consensus sequence for each barcode, we employed a read-based majority call approach. When we compared the percentage perfects obtained from each sequencing method, PacBio consistently outperformed Oxford Nanopore, with a median of 7.8% (s.d. 4.2%) higher percentage perfects, as depicted in Sup. Fig. S13. This comparison underlines the notable difference in error rates between the two sequencing platforms. It also implies that achieving comparable accuracy with Oxford Nanopore might necessitate more sophisticated consensus calling strategies, along with higher sequencing depths, to match the performance seen with PacBio.

## Conclusion

This study underscores the efficacy of Degenerate DropSynth as a robust, scalable tool for protein engineering and synthetic biology, especially in the construction of large gene libraries of chimeric proteins. We demonstrate the capability to assemble up to eight distinct variants for each gene by isolating multiple overlap-compatible fragments into emulsion PCR (ePCA) droplets, utilizing barcoded microbeads. In addition to functionally characterizing the sensor histidine kinases assembled in this study, future gene synthesis work will explore the scalability of this approach, which, we believe, has yet to reach its full potential. Potential strategies include increasing the number of variants per barcode beyond 8 while maintaining uniform degeneracy across the entire library, employing combinatorial assembly with multiple variable fragment positions, and integrating larger bead sets (exceeding 1536x) with degeneracy. Considering the observed 8% rate of perfect assemblies for genes of 1 kbp length, the use of longer lengths may require either error-correction or substantial oversampling in subsequent functional assays. In sum, Degenerate DropSynth significantly enhances the scope, length, and cost-efficiency of gene library construction, thereby facilitating the exploration and understanding of protein families through large-scale, programmable assembly.

## Acknowledgements

This work was supported by a Career Award at the Scientific Interface [to C.P.] from the Burroughs Wellcome Fund. A.S.H was supported by the NIH T32 Molecular Biology and Biophysics Training Program. We thank the UOregon GC3F core staff, Doug Turnbull and Tina Arredondo for productive discussion on the use of MAS ISO-seq. Plasmid pSR348 was a gift from Jeffrey Tabor (Addgene plasmid # 124713 ; http://n2t.net/addgene:124713 ; RRID:Addgene_124713).

## Conflicts of Interest

C.P. is a named inventor on a provisional patent based on this method.

## Methods Gene Design

All amino acid sequences (1,127,577 in total) containing a histidine kinase domain were obtained from UniProt release 2021_02. This dataset was then loaded into HMMER (version 3.2.1) and the domains were annotated for each sequence using the Pfam HMMs (version 33.1). Phobius (version 1.01) and TMHMM-py (version 1.3.1) were both used to define transmembrane regions (TMs); only TMs with an overlap consensus of at least nine residues were kept, with the boundaries determined by Phobius being used. Only proteins with 2 TM domains and a HAMP domain were kept, leaving 126,611 proteins. These were further filtered into proteins where the length from the N-terminal to the end of the HAMP domain could fit into a 4x or 5x 300-mer oligo DropSynth assembly. Sequences are provided in Supplementary Data file 1.

### DropSynth Oligo Design

The DropSynth oligos were designed using a series of custom scripts available at (https://github.com/PlesaLab/DropSynth_code_2023). These scripts were significantly optimized compared to older versions of the design scripts. Briefly some of the changes include a switch to a “recipe” based workflow with all parameters in a single file. The use of Lattice-Automation’s seqfold python library for minimum free energy structure calculations, instead of unafold (hybrid-ss-min). See Sup. Fig. S14 for a comparison. A programmable database for handling all restriction enzyme sites required. The ability to split as many genes as necessary in the first step with subsequent (384x, 1536x) library allocation. Virtual assembly and translation to verify oligo designs. The option to do microbead barcode reversal between libraries to offset barcoded microbead effects. Improved codon optimization with lower split failures through the use of several hardcoded rules. The ability to require certain sequences (controls) in each library. Single oligo processing for very small genes. Initial support for DNA (non-protein) constructs. Improved oligo junction length handling, with genes that fail due to length placed into a special file for input into higher oligo splits. As before, each for the four subpools was given a set of unique 15-mer subpool amplification primers as shown in Sup. Table S2.

### Degenerate Oligo Design

We created an R script to create all necessary degenerate oligos. Briefly we initially designed DropSynth oligos using the +2 N fusion variants, since all other variants are as long or shorter. This ensured that all variants could fit on the last oligo. All DropSynth oligos were loaded and the payload sequence between the BtsI sites was determined. The overlap sequence between the last and second to last gene fragments was determined. We then determined if the degenerate mutation could be made based on the distance between the end cloning site (GACGTGAGACC) and the end of the overlap. Since the overlap sequence could not be modified, degenerate oligos were designed only if there was sufficient length after the overlap to implement the desired mutation. If sufficient, the script made degenerate oligos by selectively removing or mutating codons for each phase variant and level of degeneracy (Fig. 1d,e). If codons were removed, the overall length of the oligo was maintained by adding random bases into the padding region between the end cloning site and assembly amplification reverse primer site (skpp504R), checking to make sure no illegal restriction sites were introduced. If codons were changed, the padding was left unchanged, but the new sequence was still screened for illegal restriction sites. All successful degenerate oligo designs were combined together with the full DropSynth oligo set. All oligo sequences are provided in Supplementary Data file 2.

### Oligo Amplification and Processing

Oligo designs were ordered as part of a pool of 58,500 300-mer oligos from Twist Bioscience. Processing followed the same protocol as detailed previously. Briefly the OLS pool was resuspended to a concentration of 19 ng/uL. Subpools were amplified using 18 - 20 cycles (as first determined using qPCR) with Kapa HiFi. Bulk amplification of each subpool was then carried out using 8 - 11 cycles (determined by qPCR) with 0.5 to 1 ng of template. Between 7 - 9 ug of bulk amplified DNA was put into the nt.BspQI nick processing, with a yield ranging from 4.2 - 6.2 ug (corresponding to a range of molar yields of 52 - 75%).

### Emulsion Assembly and Suppression PCR

Briefly, the DropSynth reaction was carried out using 1.3 ug of processed DNA. After emulsion breaking with chloroform, the correct length assemblies were (blind) size selected with an agarose gel. DNA was extracted from gel slices with an NEB Monarch DNA Gel Extraction Kit and eluted in 30 uL. Of this 1 uL was used as template in a suppression PCR reaction (first on a qPCR) using 25 - 28 cycles. These reactions were cleaned up, quantified by Qubit, and 0.36 pmol was subsequently used in each Golden Gate reaction as described below.

### Plasmid Design of pHKGG1

Plasmid pSR348 containing the complete NarX SHK under the LacI inducible promoter was received from Addgene (#124713) and sequence verified using full plasmid nanopore sequencing. Golden Gate Assembly was used to change the antibiotic selection maker from spectinomycin resistance to carbenicillin creating the plasmid pSR348_Carb. Several clones were isolated and their plasmids extracted and sequence verified in the same manner as pSR348. After swapping the selection marker, we replaced the NarX CDS with the wild type EnvZ SHK (amplified from E. *coli* MG1655) to generate the plasmid EnvZ_pSR348_Carb. Finally, site directed mutagenesis was used to remove a KpnI restriction site for backup cloning purposes creating the final plasmid renamed pHKGG1. A complete plasmid map and all primers used to generate pHKGG1 are found in the supporting information (Sup. Fig. S15) and Sup. Table S1. All transformations after each cloning step were done in 10-Beta cells (NEB) unless otherwise noted.

### Golden Gate Assembly

A three fragment Golden Gate assembly was used to clone our DropSynth generated libraries. The first fragment (A) of the assembly consisted of a variable segment composed of HK sensory domains produced from the degenerate DropySynth protocol. The overhang CATA was used at the start, where the final adenine base is the start of the ATG codon of the gene. The overhang GACG was used at the end where GAC encodes residue D232 in the EnvZ protein. The second fragment (B) contained a conserved portion of the EnvZ gene corresponding to amino acid residues D232-G450 and a 24 basepair quasi-randomer barcode region (NNBBDDBBVVHHDDBBVVHNDDNN) downstream of the stop codon. The final fragment (C) was derived from pHKGG1 and contained the p15A Ori, Amp selection marker, lac repressor, and P_trc_ inducible promoter. The fragment (B) containing the conserved portion of EnvZ and the 24 bp barcode along with pHKGG1 backbone were both generated via PCR and their respective primers are tabulated in the Sup. Table S1. The sequences of the three Golden Gate fragments are provided in Sup. Table S4.

For each Degenerate DropSynth library four Golden Gate reactions were run using the following temperature program: 1) Incubate at 37°C for 20 hours 2) heat inactivation of both enzymes at 80°C for 20 mins 3) final hold at 12°C. The molar ratios used were 0.18 pmol for FrgC (backbone), 0.36 pmol for FrgB (conserved region of EnvZ and barcode), and 0.36 pmol for FrgA (degenerate DropSynth synthesized library) for a total of 470 ng of DNA in the reaction. The reagent volumes were 2.5 uL of 10x T4 DNA Ligase Buffer (NEB), 2.5 uL of 10 mM ATP (NEB), 0.25 uL T4 DNA Ligase (NEB), 0.75 uL BsaI-HFv2 (NEB), and water to complete the remaining volume to a total of 25 uL. Assemblies were then pooled and cleaned using the Monarch PCR and DNA Cleanup Kit (NEB) and then drop dialyzed on a 0.05 um membrane filter (Sigma Millipore) for a minimum of 30 minutes. Purified DNA was used as the input for two electroporations (Bio-Rad MicroPulser) which were then combined and plated. Serial dilutions were used to calculate the total number of CFUs and are listed in Sup. Table S3.

### Assembly Barcode Sequencing (MAS ISO-seq) and Analysis

Each library was PCR amplified with one of the HK_PB_0#_FWD+REV primer pairs using Q5 DNA polymerase, 10 ng of template (miniprepped plasmid), in a 50uL reaction for 11-12 cycles. To submit samples for MAS ISO-seq, the four libraries were mixed into a single 30μL sample containing 1.25ng/μL per library (5ng/μL total) and submitted to the UOregon GC3F core for library preparation and sequencing. Briefly, PacBio MAS ISO-Seq for 10x Single Cell 3’ kit (102-659-600) was used to generate arrays for sequencing on a Sequel II instrument producing 120.20 Gbases of unique molecular data in 6.14 million raw reads. These samples were also sequenced by Plasmidsaurus using Oxford Nanopore sequencing with R10.4.1 flowcells, v14 chemistry, and basecalled with Guppy 6.5.7. This produced between 10,592,215 to 12,245,627 reads for each sample. All sequencing data are available from the NIH sequencing read archive (SRA) under the BioProject PRJNA1049019.

For the MAS ISO-seq data, skera (0.1.0) was used to split the MAS arrays within 1,268,200 CCS reads, producing 15,907,686 split segments. Lima was then used to demultiplex the split segments resulting in 2,740,434 to 4,826,815 reads per library. For both MAS ISO-seq and Oxford Nanopore data a custom python script was used to first identify the constant regions flanking the barcode (GTCGCTGCCGAACAGC-24N-AGGAGAAGAGCGCACG) allowing up to 3 mismatches. Then each read was scanned for the presence of the NdeI (CATATG) site at the start codon and the conserved region in the vector immediately flanking the cloning site (GACGACCGCACGCTGCTG which corresponds to residues 232-237 (DDRTLL) of envZ). The script outputs this variable region and each associated barcode, as well as the barcodes counts. Barcode counts were input into starcode (1.4) for collapse with a distance of 1 using the sphere algorithm. A consensus call was made for each barcode using a simple majority call. Genes were aligned to their closest parent sequence using minimap2 with k-mers set to 10. To assign reads to degenerate variants the last 16 bp of each variable region was taken. Perfect matches to designs were taken as is, for the remainder we calculated the Levenshtein distance between each 16 bp end sequence and all degenerate variants for that parent sequence. The read was assigned based on the smallest Levenshtein distance. In case of a tie (rare), a random assignment was made. All subsequent analysis and plots were carried out in R (4.3.1).

### Error Model

The percentage of perfect oligos from oligo synthesis are given by

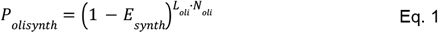

where L_oli_ is the length of the oligos (300 nt), N_oli_ is the number of oligos used in the assembly (4 or 5), and E_synth_ is the estimated error rate of the synthesis process, discussed in text. The errors introduced by PCR amplification can be split into the amplification of the oligos and the suppression PCR of the assembled genes. The percentage of perfects after both PCRs should be given by

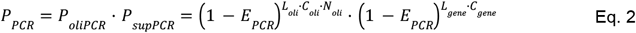

where L_gene_ is the length of the gene, C_oli_ is the number of PCR cycles used in oligos amplification, C_gene_ is the number of suppression PCR cycles used in gene amplification, and E_PCR_ is the error rate of the polymerase, discussed in text. We model the ePCA process using

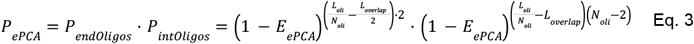

where the first part accounts for the two oligos on the end while the latter part accounts for internal oligos. L_overlap_ is the typical length of the overlap between fragments and E_ePCA_ is the error rate of the assembly process. The total estimated number of percentage perfects is then given by

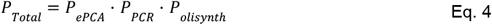

The only unknown parameter which is allowed to vary during fits is E_ePCA_.

## Notes

https://www.ncbi.nlm.nih.gov/bioproject/?term=PRJNA1049019

https://github.com/PlesaLab/DropSynth_code_2023

https://dropsynth.org/

### Competing Interest Statement

A provisional patent has been filed related to this work.

## References

(1) Yu, K.; Liu, C.; Kim, B.-G.; Lee, D.-Y. Synthetic Fusion Protein Design and Applications. Biotechnol. Adv. 2015, 33 (1), 155–164.

(2) Lin, C.-Y.; Liu, J. C. Modular Protein Domains: An Engineering Approach toward Functional Biomaterials. Curr. Opin. Biotechnol. 2016, 40, 56–63.

(3) Koide, S. Generation of New Protein Functions by Nonhomologous Combinations and Rearrangements of Domains and Modules. Curr. Opin. Biotechnol. 2009, 20 (4), 398–404.

(4) Maervoet, V. E. T.; Briers, Y. Synthetic Biology of Modular Proteins. Bioengineered 2017, 8 (3), 196–202.

(5) Gordley, R. M.; Bugaj, L. J.; Lim, W. A. Modular Engineering of Cellular Signaling Proteins and Networks. Curr. Opin. Struct. Biol. 2016, 39, 106–114.

(6) Punt, P. J.; Levasseur, A.; Visser, H.; Wery, J.; Record, E. Fungal Protein Production: Design and Production of Chimeric Proteins. Annu. Rev. Microbiol. 2011, 65, 57–69.

(7) Menzella, H. G.; Reeves, C. D. Combinatorial Biosynthesis for Drug Development. Curr. Opin. Microbiol. 2007, 10 (3), 238–245.

(8) Vymětal, J.; Mertová, K.; Boušová, K.; Šulc, J.; Tripsianes, K.; Vondrasek, J. Fusion of Two Unrelated Protein Domains in a Chimera Protein and Its 3D Prediction: Justification of the X-Ray Reference Structures as a Prediction Benchmark. Proteins 2022, 90 (12), 2067–2079.

(9) Patel, D. K.; Menon, D. V.; Patel, D. H.; Dave, G. Linkers: A Synergistic Way for the Synthesis of Chimeric Proteins. Protein Expr. Purif. 2022, 191, 106012.

(10) Klein, J. S.; Jiang, S.; Galimidi, R. P.; Keeffe, J. R.; Bjorkman, P. J. Design and Characterization of Structured Protein Linkers with Differing Flexibilities. Protein Eng. Des. Sel. 2014, 27 (10), 325–330.

(11) Nielsen, M. L.; Isbrandt, T.; Petersen, L. M.; Mortensen, U. H.; Andersen, M. R.; Hoof, J. B.; Larsen, T. O. Linker Flexibility Facilitates Module Exchange in Fungal Hybrid PKS-NRPS Engineering. PLoS One 2016, 11 (8), e0161199.

(12) Sieber, V.; Martinez, C. A.; Arnold, F. H. Libraries of Hybrid Proteins from Distantly Related Sequences. Nat. Biotechnol. 2001, 19 (5), 456–460.

(13) Ostermeier, M.; Shim, J. H.; Benkovic, S. J. A Combinatorial Approach to Hybrid Enzymes Independent of DNA Homology. Nat. Biotechnol. 1999, 17 (12), 1205–1209.

(14) Ohlendorf, R.; Schumacher, C. H.; Richter, F.; Möglich, A. Library-Aided Probing of Linker Determinants in Hybrid Photoreceptors. ACS Synth. Biol. 2016, 5 (10), 1117–1126.

(15) Sidore, A. M.; Plesa, C.; Samson, J. A.; Lubock, N. B.; Kosuri, S. DropSynth 2.0: High-Fidelity Multiplexed Gene Synthesis in Emulsions. Nucleic Acids Res. 2020, 740977.

(16) Borovkov, A. Y.; Loskutov, A. V.; Robida, M. D.; Day, K. M.; Cano, J. A.; Le Olson, T.; Patel, H.; Brown, K.; Hunter, P. D.; Sykes, K. F. High-Quality Gene Assembly Directly from Unpurified Mixtures of Microarray-Synthesized Oligonucleotides. Nucleic Acids Res. 2010, 38 (19), e180.

(17) Chakraborty, S.; Li, M.; Chatterjee, C.; Sivaraman, J.; Leung, K. Y.; Mok, Y.-K. Temperature and Mg2+ Sensing by a Novel PhoP-PhoQ Two-Component System for Regulation of Virulence in Edwardsiella Tarda. J. Biol. Chem. 2010, 285 (50), 38876–38888.

(18) Matilla, M. A.; Velando, F.; Martín-Mora, D.; Monteagudo-Cascales, E.; Krell, T. A Catalogue of Signal Molecules That Interact with Sensor Kinases, Chemoreceptors and Transcriptional Regulators. FEMS Microbiol. Rev. 2022, 46 (1). 10.1093/femsre/fuab043.

(19) Hirose, Y.; Shimada, T.; Narikawa, R.; Katayama, M.; Ikeuchi, M. Cyanobacteriochrome CcaS Is the Green Light Receptor That Induces the Expression of Phycobilisome Linker Protein. Proc. Natl. Acad. Sci. U. S. A. 2008, 105 (28), 9528–9533.

(20) Xu, Y.; Zhao, Z.; Tong, W.; Ding, Y.; Liu, B.; Shi, Y.; Wang, J.; Sun, S.; Liu, M.; Wang, Y.; Qi, Q.; Xian, M.; Zhao, G. An Acid-Tolerance Response System Protecting Exponentially Growing Escherichia Coli. Nat. Commun. 2020, 11 (1), 1496.

(21) Cai, S. J.; Inouye, M. EnvZ-OmpR Interaction and Osmoregulation in Escherichia Coli. J. Biol. Chem. 2002, 277 (27), 24155–24161.

(22) Bhate, M. P.; Molnar, K. S.; Goulian, M.; DeGrado, W. F. Signal Transduction in Histidine Kinases: Insights from New Structures. Structure 2015, 23 (6), 981–994.

(23) Buschiazzo, A.; Trajtenberg, F. Two-Component Sensing and Regulation: How Do Histidine Kinases Talk with Response Regulators at the Molecular Level? Annu. Rev. Microbiol. 2019, 73, 507–528.

(24) Jung, K.; Fabiani, F.; Hoyer, E.; Lassak, J. Bacterial Transmembrane Signalling Systems and Their Engineering for Biosensing. Open Biol. 2018, 8 (4). 10.1098/rsob.180023.

(25) Bi, S.; Pollard, A. M.; Yang, Y.; Jin, F.; Sourjik, V. Engineering Hybrid Chemotaxis Receptors in Bacteria. ACS Synth. Biol. 2016, 5 (9), 989–1001.

(26) Ganesh, I.; Kim, T. W.; Na, J.-G.; Eom, G. T.; Hong, S. H. Engineering Escherichia Coli to Sense Non-Native Environmental Stimuli: Synthetic Chimera Two-Component Systems. Biotechnol. Bioprocess Eng. 2019, 24 (1), 12–22.

(27) Wang, B.; Barahona, M.; Buck, M.; Schumacher, J. Rewiring Cell Signalling through Chimaeric Regulatory Protein Engineering. Biochem. Soc. Trans. 2013, 41 (5), 1195–1200.

(28) Schmidl, S. R.; Sheth, R. U.; Wu, A.; Tabor, J. J. Refactoring and Optimization of Light-Switchable Escherichia Coli Two-Component Systems. ACS Synth. Biol. 2014, 3 (11), 820–831.

(29) Lazar, J. T.; Tabor, J. J. Bacterial Two-Component Systems as Sensors for Synthetic Biology Applications. Curr Opin Syst Biol 2021, 28. 10.1016/j.coisb.2021.100398.

(30) Wang, B.; Zhao, A.; Novick, R. P.; Muir, T. W. Activation and Inhibition of the Receptor Histidine Kinase AgrC Occurs through Opposite Helical Transduction Motions. Mol. Cell 2014, 53 (6), 929–940.

(31) Kaur, H.; Singh, S.; Rathore, Y. S.; Sharma, A.; Furukawa, K.; Hohmann, S.; Ashish Mondal, A. K. Differential Role of HAMP-like Linkers in Regulating the Functionality of the Group III Histidine Kinase DhNik1p. J. Biol. Chem. 2014, 289 (29), 20245–20258.

(32) Parkinson, J. S. Signaling Mechanisms of HAMP Domains in Chemoreceptors and Sensor Kinases. Annu. Rev. Microbiol. 2010, 64, 101–122.

(33) Stewart, V.; Chen, L.-L. The S Helix Mediates Signal Transmission as a HAMP Domain Coiled-Coil Extension in the NarX Nitrate Sensor from Escherichia Coli K-12. J. Bacteriol. 2010, 192 (3), 734–745.

(34) Airola, M. V.; Watts, K. J.; Bilwes, A. M.; Crane, B. R. Structure of Concatenated HAMP Domains Provides a Mechanism for Signal Transduction. Structure 2010, 18 (4), 436–448.

(35) Schmidl, S. R.; Ekness, F.; Sofjan, K.; Daeffler, K. N.-M.; Brink, K. R.; Landry, B. P.; Gerhardt, K. P.; Dyulgyarov, N.; Sheth, R. U.; Tabor, J. J. Rewiring Bacterial Two-Component Systems by Modular DNA-Binding Domain Swapping. Nat. Chem. Biol. 2019, 15 (7), 690–698.

(36) Al’Khafaji, A. M.; Smith, J. T.; Garimella, K. V.; Babadi, M.; Popic, V.; Sade-Feldman, M.; Gatzen, M.; Sarkizova, S.; Schwartz, M. A.; Blaum, E. M.; Day, A.; Costello, M.; Bowers, T.; Gabriel, S.; Banks, E.; Philippakis, A. A.; Boland, G. M.; Blainey, P. C.; Hacohen, N. High-Throughput RNA Isoform Sequencing Using Programmed cDNA Concatenation. Nat. Biotechnol. 2023. 10.1038/s41587-023-01815-7.

(37) Kosuri, S.; Church, G. M. Large-Scale de Novo DNA Synthesis: Technologies and Applications. Nat. Methods 2014, 11 (5), 499–507.

(38) Li, H. Minimap2: Pairwise Alignment for Nucleotide Sequences. Bioinformatics 2018, 34 (18), 3094–3100.

(39) Needleman, S. B.; Wunsch, C. D. A General Method Applicable to the Search for Similarities in the Amino Acid Sequence of Two Proteins. J. Mol. Biol. 1970, 48 (3), 443–453.

(40) Hestand, M. S.; Van Houdt, J.; Cristofoli, F.; Vermeesch, J. R. Polymerase Specific Error Rates and Profiles Identified by Single Molecule Sequencing. Mutat. Res. 2016, 784-785, 39–45.

(41) Karst, S. M.; Ziels, R. M.; Kirkegaard, R. H.; Sørensen, E. A.; McDonald, D.; Zhu, Q.; Knight, R.; Albertsen, M. High-Accuracy Long-Read Amplicon Sequences Using Unique Molecular Identifiers with Nanopore or PacBio Sequencing. Nat. Methods 2021, 18 (2), 165–169.

(42) Zurek, P. J.; Knyphausen, P.; Neufeld, K.; Pushpanath, A.; Hollfelder, F. UMI-Linked Consensus Sequencing Enables Phylogenetic Analysis of Directed Evolution. Nat. Commun. 2020, 11 (1), 6023.

