## Supplementary Information for "Degenerate DropSynth for Simultaneous Assembly of Diverse Gene Libraries and Local Designed Mutants"

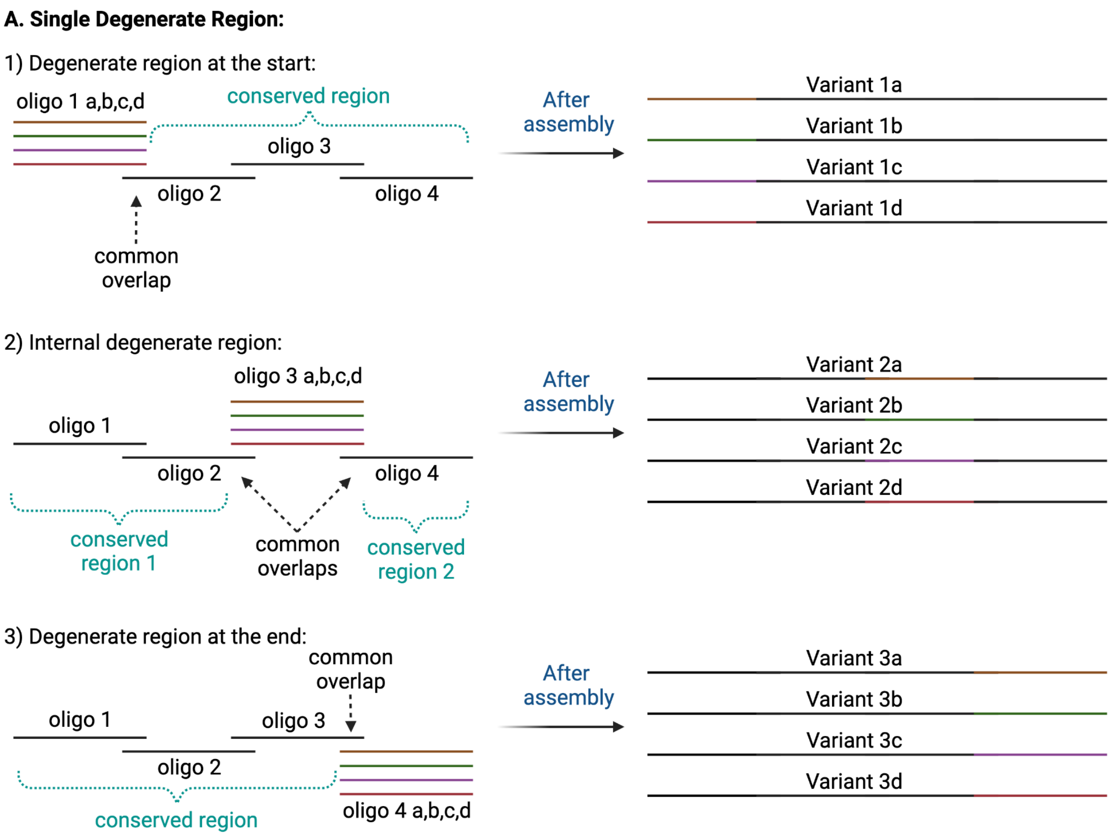


**Supplementary Figure S1** - Degenerate DropSynth could be used to introduce variation in many parts of an assembled gene by creating multiple oligos for gene fragments at **1)** the start, **2)** internally, or **3)** at the end.

**B. Multiple Degenerate Regions (Combinatorial):**

4) Combinatorial example with 2 regions, (start and internal):

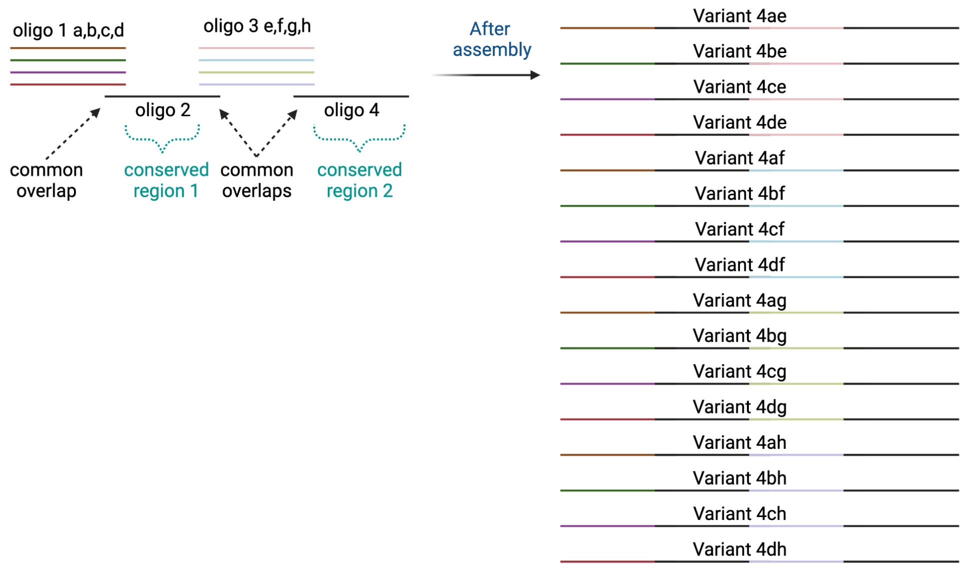


**Supplementary Figure S2** - Introducing degeneracy in multiple fragments leads to combinatorial assemblies.

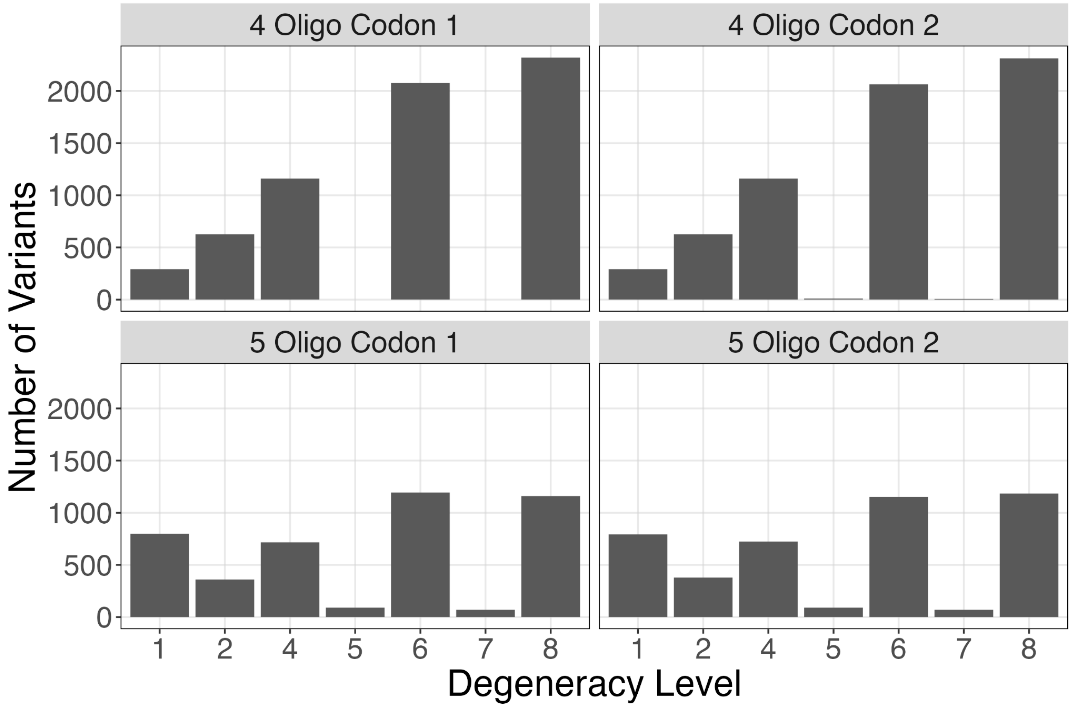


**Supplementary Figure S3** - The absolute number of variants designed at each degeneracy level for each of the four libraries tested.

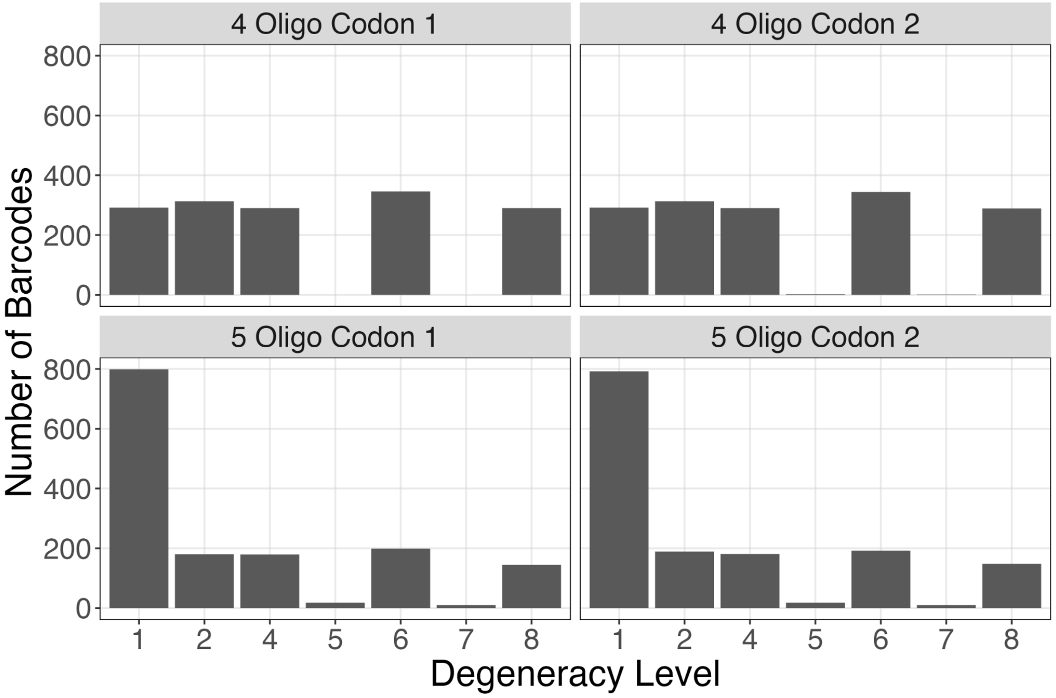


**Supplementary Figure S4** - The number of barcodes used (out of 1536) for each degeneracy level in each of the four libraries tested.

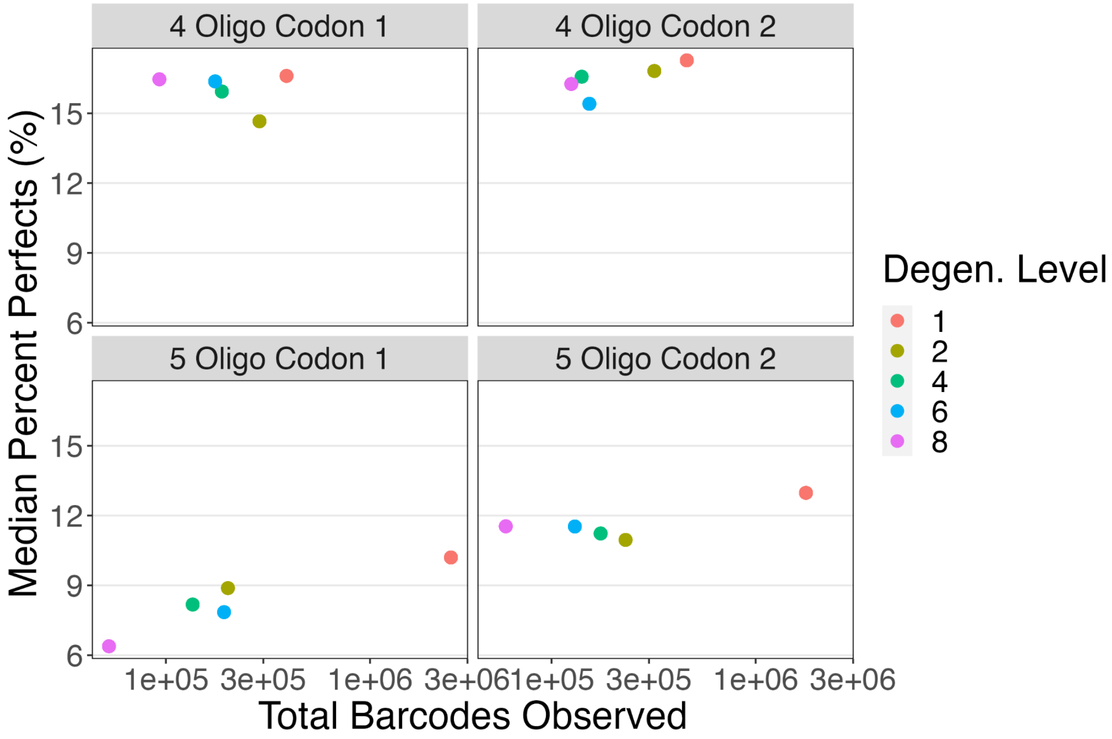


**Supplementary Figure S5** - When plotting the median percentage of perfects, we note a correlation with the total number of barcodes observed and the degeneracy levels which is much stronger with the 5 oligo libraries (0.88 and 0.96 Pearson) (bottom-row) compared to the 4 oligo libraries (-0.38 and 0.67 Pearson) (top-row).

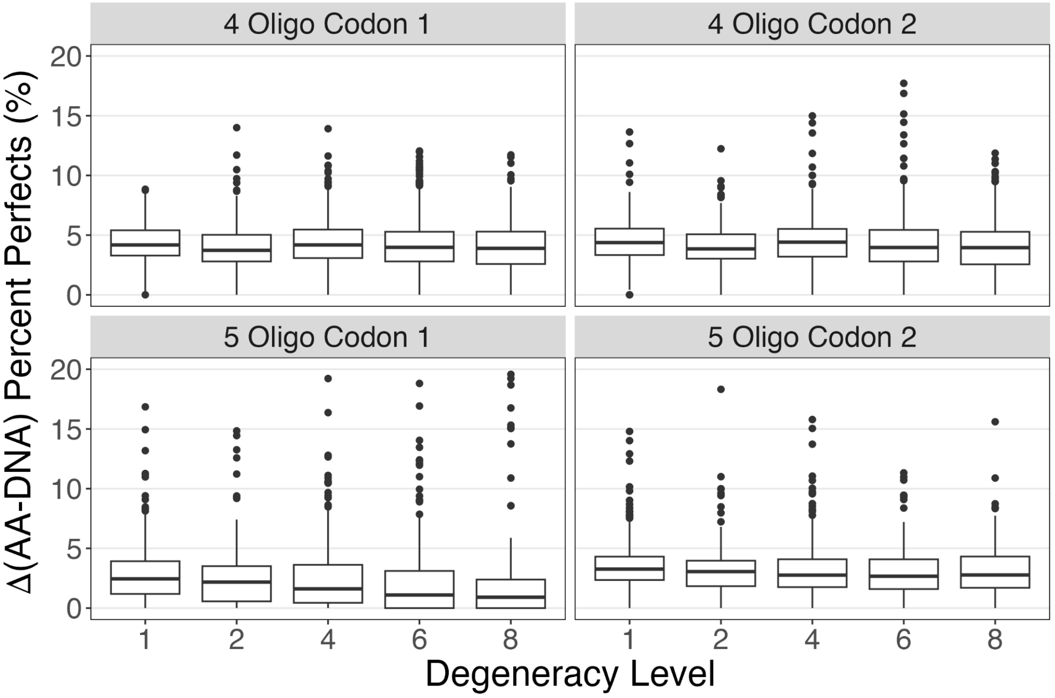


**Supplementary Figure S6** - The difference in percentage perfects observed at the protein amino acid level (synonymous mutations collapsed) and the DNA level. We see a relatively consistent difference of 4.0% (s.d. 0.2%) with 4 oligos and 2.7% (s.d. 0.8%) with 5 oligos.

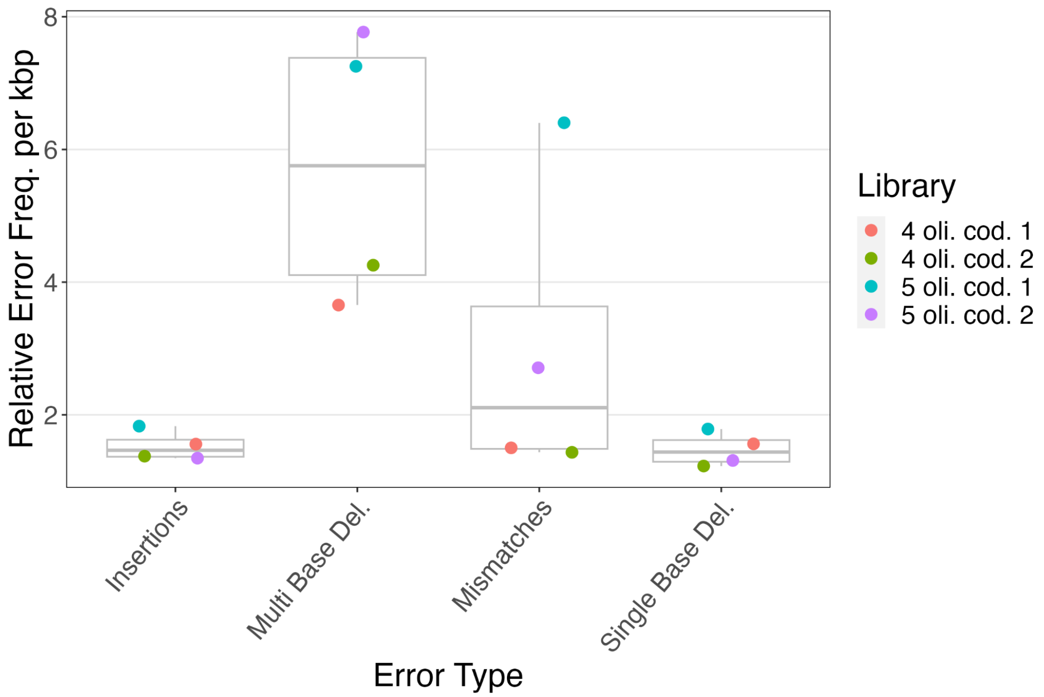


**Supplementary Figure S7** - Analysis of the CIGAR alignment strings produced by minimap2 reveals the relative error frequencies per kbp for different types of errors. We find equivalent rates of insertions and single base deletions among the four libraries and much higher rates of multi base deletions. In the 4-oligo libraries we see mismatch rates comparable to insertions and single base deletions, while in the 5-oligo libraries mismatches are substantially higher.

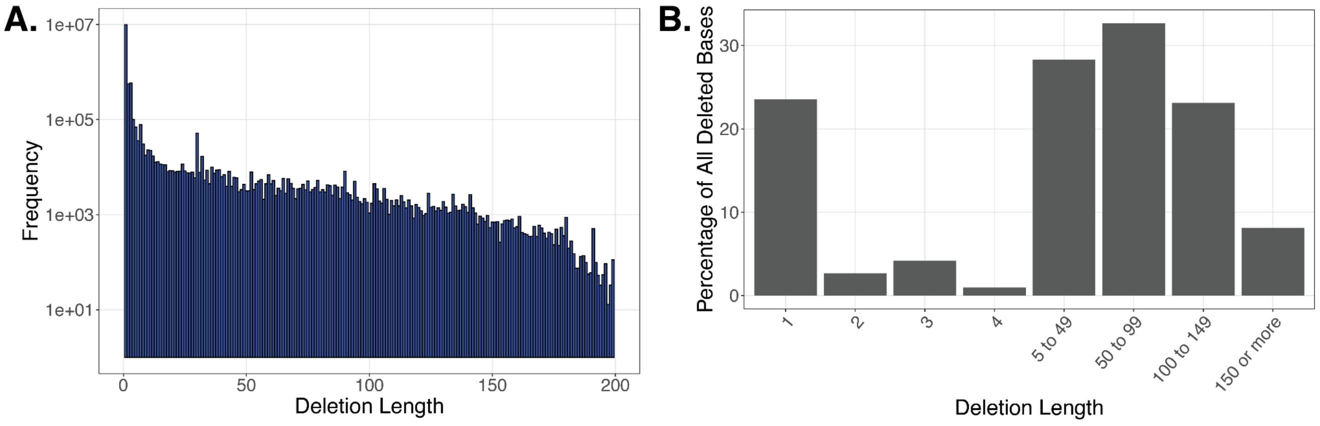


**Supplementary Figure S8 - A.** A histogram of the length of deletions observed across all 4 libraires. While single-base deletions are dominant a substantial amount of long deletions are observed. **B.** Since many multi-base deletions are so long. At any given base in a deletion, only 23.6% are from single-deletions, while 28.3% are deletion of 5-49 bp in length, 32.7% are deletions of 50 to 99 bp in length, 23.1% are deletions of 100 to 149 bp in length, and 8.1% are 150 bp or longer.

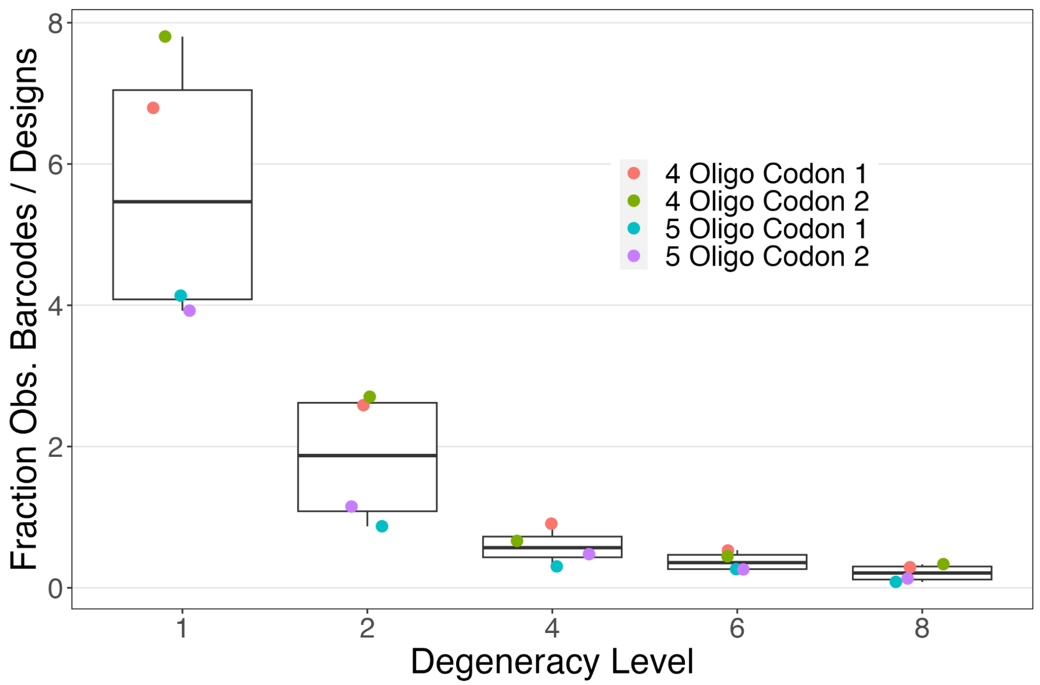


**Supplementary Figure S9 -** The fraction of barcodes observed, normalized by the total faction of designs. For each library, we determined the sum of all observed unique gene barcodes at a particular degeneracy level, and divided it by the total number of observed unique gene barcodes in the library, to calculate the fraction of observed barcodes. We determined the fraction of designs at each degeneracy level by dividing the total number of variants at that degeneracy level divided by the total number of variants in the entire library. We see a strong decay in the observed barcodes as the degeneracy level is increased.

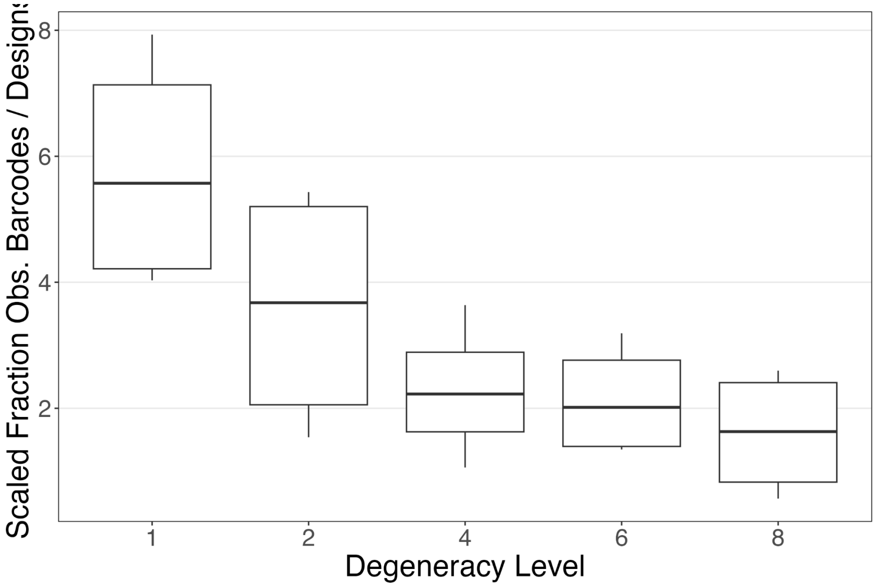


**Supplementary Figure S10** - The fraction of barcodes observed, divided by the total faction of designs, scaled by degeneracy level. In other words we take the data from Supplementary Figure S9 on the fraction of observed barcodes and we multiply it by the degeneracy level. If we assume the amount of DNA for variants at the end of assembly is inversely proportional to the degeneracy level we would expect roughly similar numbers (eg. a variant from a degeneracy level of 8 has ⅛ the amount of DNA as a variant from degeneracy level of 1). Since we still see a strong decay, this implies that other factors such as PCR amplification contribute to the effect.

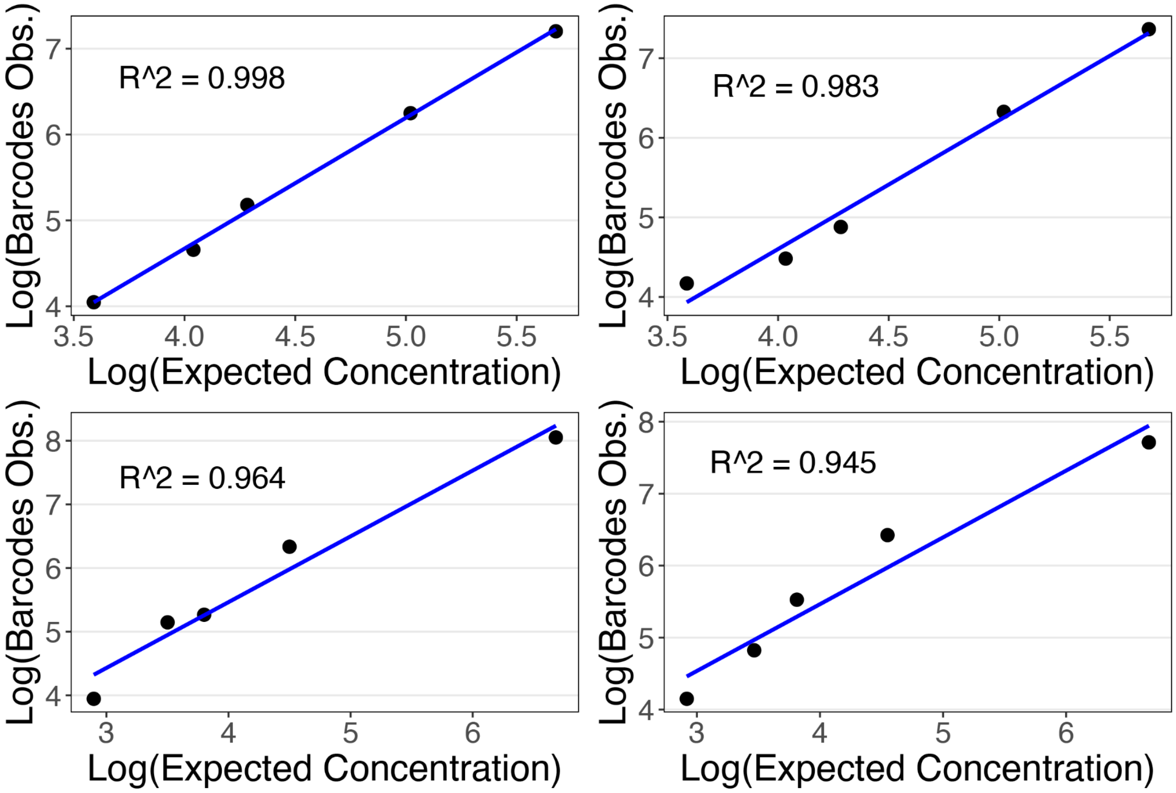


**Supplementary Figure S11** - A model of PCR amplification applied to all four libraries. The y-axis values are log transformed barcodes observed per variant while the x-axis is the expected variant concentration given by the total number of barcoded beads with a given degeneracy level divided by the total number of variants at that level.

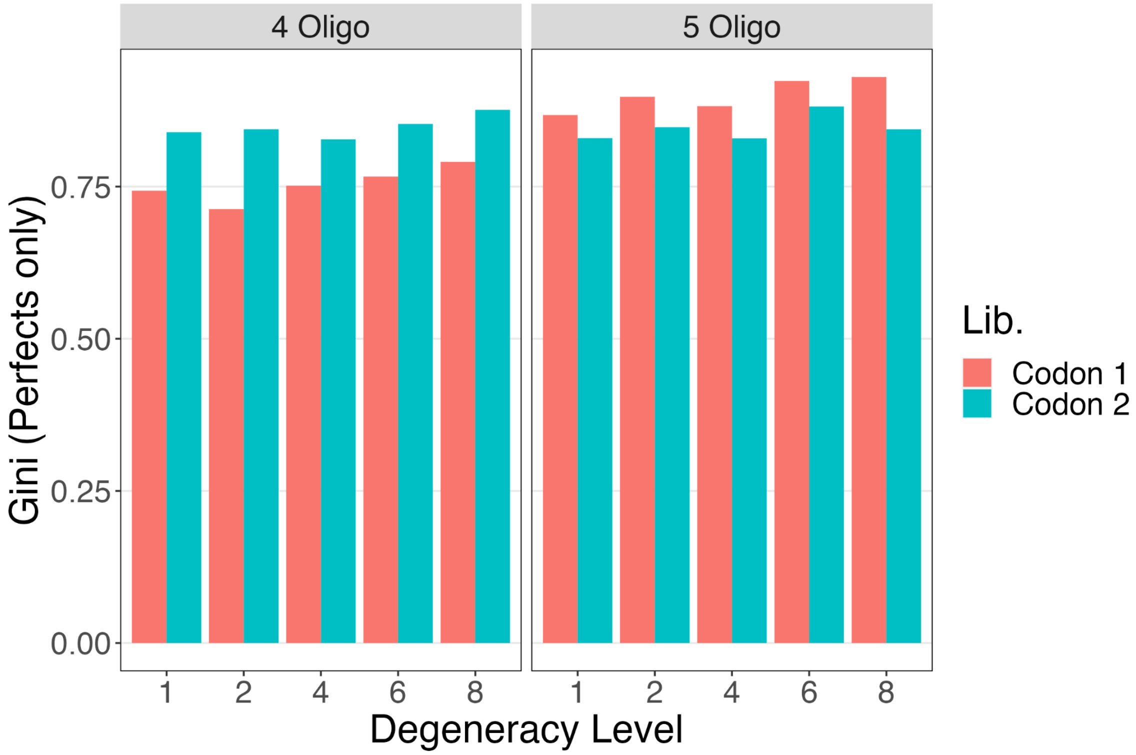


**Supplementary Figure S12** - The Gini coefficient is a measure of the inequality among the distribution of library members. The values observed are consistent with previous libraries assembled with DropSynth.

A.

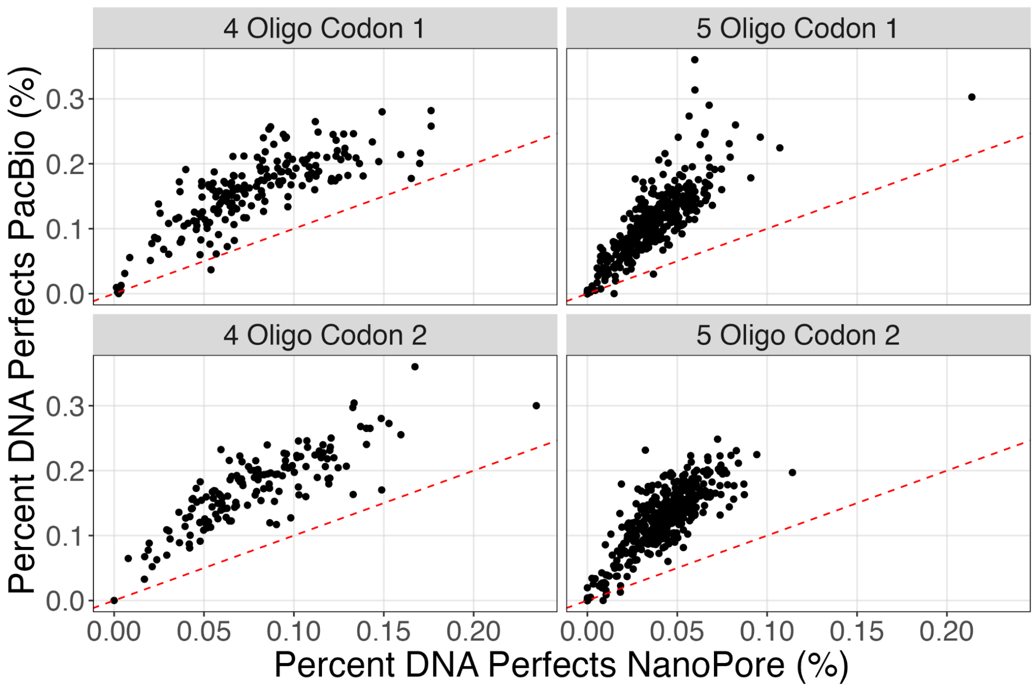


B.

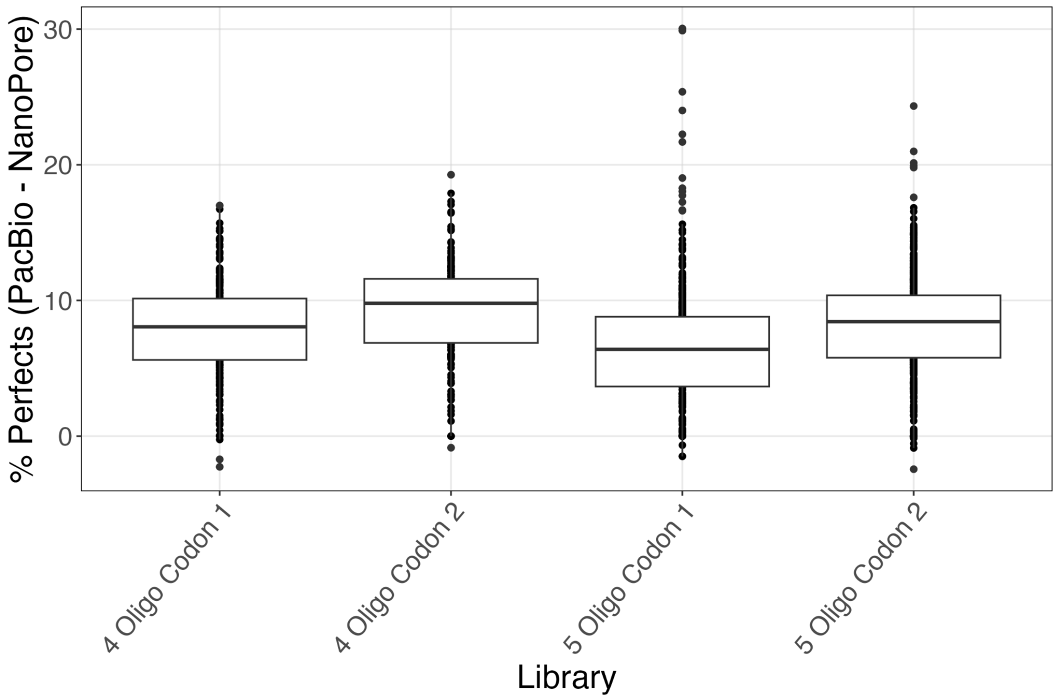


**Supplementary Figure S13 - A.** The percentage of perfects determined from the PacBio data (y-axis) plotted against the percentage of perfects determined using Oxford Nanopore data (x-axis). Data using only degeneracy level of 1. **B.** The distribution of delta percentage perfects shows a consistent (median) 7.8% higher rate for PacBio data highlighting its lower error rate in sequencing.

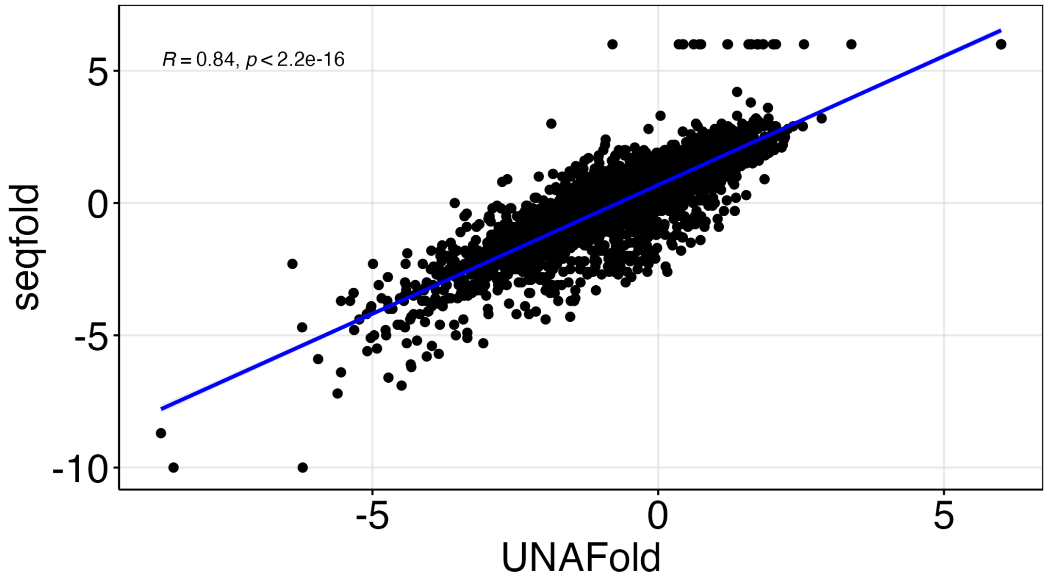


**Supplementary Figure S14** - The folding energy of 4000 random 20 bp sequences determined using both seqfold and UNAfold, shows a 0.84 correlation.

**Supplementary Table S1 -** Primer sequences used in this study.

| Oligo Name | Sequence (annealing region highlighted) | Purpose |
|---|---|---|
| pSEVA121_AB_CARB_FWD | GAGAACGGTCTCCgtaaattagtagcccgcctaa | pSR348 → pSR348_Carb |
| pSEVA121_AB_CARB_REV | GAGAACGGTCTCCgggtcgtccaaaaaaaaagg | |
| pSR348_AB_CARB_FWD | GAGAACGGTCTCCaccccccagtatcagcccgtca | |
| pSR348_AB_CARB_REV | GAGAACGGTCTCCttacgaaacgatcctcatcc | |
| pSR348_AB_CARB_FWD | GAGAACGGTCTCCaccccccagtatcagcccgtca | pSR348_Carb → EnvZ_pSR348_Carb |
| pSR348_AB_CARB_REV | GAGAACGGTCTCCttacgaaacgatcctcatcc | |
| EnvZ_WT_FWD | GTCATCGGTCTCCacatgAGGCGATTGCGCTTC | |
| EnvZ_WT_REV | GTCATCGGTCTCCttaCCCTTCTTTTGTCGTGC | |
| pSR348_delKpnI_FWD | aggggatcctctagagtcgac | KpnI site directed mutagenesis |
| pSR348_delKpnI_REV | gtacctataaacgcagaaaggc | |
| FrgC_HKBC2_FWD | CACCTCGGTCTcAAGAAGAGCgcacGACGTcaCgtCgcaGAATTCcttttcggggaaatgtgcgcggaacccgggtaataataatctagaccaggcatc | Make Fragment C |
| FrgC_HKBC2_REV | GGCACTGGTCTcatAtgagttgtgcgatttaaattaggagag | |
| frgB_bH_envZ_FWD1 | gcCACCTCGGTCTcggaCgaccgcacgctgctga | Make Fragment B - PCR 1 |
| frgB_bH_envZ_REV1 | gctGTTCgGCAGCgACACttaGGTACCTTATTGGGAGGTttacccttcttttgtcgtgc | |
| frgB_bH_envZ_FWD2 | gcCACCTCGGTCTcggaC | Make Fragment B - PCR 2 - Add barcode |
| frgB_bH_envZ_REV2 | tgcGCTGGTCTcCTTCTcctNNHHNDBBVVHHDDBBVVHHVVNNgctGTTCgGCAGCgACAC | |
| HK_PB_02_FWD | CTACACGACGCTCTTCCGATCTACACACAGACTGTGAGCACACAGCACTCTCCTAATTTAAATCGCACAACTCATATG | PacBio lib. prep. |
| HK_PB_02_REV | AAGCAGTGGTATCAACGCAGAGCTCACAGTCTGTGTGTCGTGACGTCGTGCGCTCTTCT | |
| HK_PB_03_FWD | CTACACGACGCTCTTCCGATCTACACATCTCGTGAGAGCACACAGCACTCTCCTAATTTAAATCGCACAACTCATATG | |
| HK_PB_03_REV | AAGCAGTGGTATCAACGCAGAGCTCTCACGAGATGTGTCGTGACGTCGTGCGCTCTTCT | |
| HK_PB_06_FWD | CTACACGACGCTCTTCCGATCTCATATATATCAGCTGTCACACAGCACTCTCCTAATTTAAATCGCACAACTCATATG | |
| HK_PB_06_REV | AAGCAGTGGTATCAACGCAGAGACAGCTGATATATATGCGTGACGTCGTGCGCTCTTCT | |
| HK_PB_07_FWD | CTACACGACGCTCTTCCGATCTTCTGTATCTCTATGTGCACACAGCACTCTCCTAATTTAAATCGCACAACTCATATG | |
| HK_PB_07_REV | AAGCAGTGGTATCAACGCAGAGCACATAGAGATACAGACGTGACGTCGTGCGCTCTTCT | |

**Supplementary Table S2 -** The subpool amplification primers.

| Library | Oligo Name | Sequence |
|---|---|---|
| 4 Oligo Codon 1 | skpp15-9-F filt15-453 | Biotin-CGATCGTGCCCACCT |
| 4 Oligo Codon 1 | skpp15-9-R filt15-1189 | GTGCGGGCTCCAACT |
| 4 Oligo Codon 2 | skpp15-13-F filt15-286 | Biotin-GGGTTCGAGCGGGAG |
| 4 Oligo Codon 2 | skpp15-13-R filt15-11 | TAGCGCGCAGAGAGG |
| 5 Oligo Codon 1 | skpp15-26-F filt15-16376 | Biotin-GCGGCACCACAAACT |
| 5 Oligo Codon 1 | skpp15-26-R filt15-327 | CGTGGCCTCTGTCCT |
| 5 Oligo Codon 2 | skpp15-28-F filt15-295 | Biotin-GACTGCGGCGTTGGT |
| 5 Oligo Codon 2 | skpp15-28-R filt15-2129 | TACGCCCGGGACAGA |

**Supplementary Table S3 -** The number of CFUs observed after transformation.

| Library | Total CFUs |
|---|---|
| 4 Oligo Codon 1 | 2.49 x 10^6 |
| 4 Oligo Codon 2 | 2.1 x 10^6 |
| 5 Oligo Codon 1 | 14.1 x 10^6 |
| 5 Oligo Codon 2 | 66 x 10^6 |

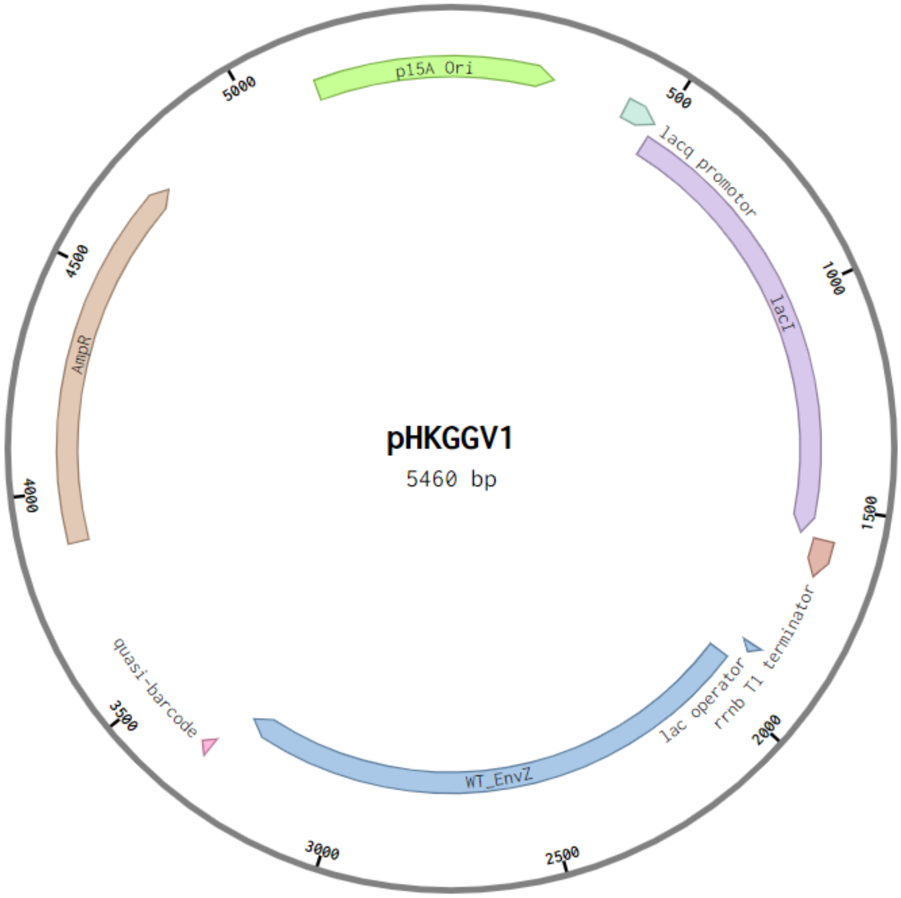


**Supplementary Figure S15** - The map of plasmid pHKGGV1. Our libraries are cloned into the N-terminal region of EnvZ. This plasmid is a derivative of plasmid pSR348 from Dr. Jeffrey Tabor's lab.

**Supplementary Table S4 -** The sequences of the three fragments used in Golden Gate to make libraries in plasmid pHKGGV1. The BsaI-HF-V2 recognition seq is shown in blue and overhang seq in red.

| Fragment | Sequence |
|---|---|
| **Frg_A - variable region** | GGCACTGGTCTcacatatg NNNNNNNNNN gacgtgAGACCGAGGTGgc |
| **Frg_B - C terminal portion of envZ and barcode** | gcCACCTCGGTCTcggaCgaccgcacgctgctgatggcgggggtaagtcacgacttgcgcacgccgct<br>gacgcgtattcgcctggcgactgagatgatgagcgagcaggatggctatctggcagaatcgatcaataaagata<br>tcgaagagtgcaacgccatcattgagcagtttatcgactacctgcgcaccgggcaggagatgccgatggaaat<br>ggcggatcttaatgcagtactcggtgaggtgattgctgccgaaagtggctatgagcgggaaattgaaaccgcgct<br>ttaccccggcagcattgaagtgaaaatgcacccgctgtcgatcaaacgcgcggtggcgaatatggtggtcaacg<br>ccgcccgttatggcaatggctggatcaaagtcagcagcggaacggagccgaatcgcgcctggttccaggtgga<br>agatgacggtccgggaattgcgccggaacaacgtaagcacctgttccagccgtttgtccgcggcgacagtgcg<br>cgcaccattagcggcacgggattagggctggcaattgtgcagcgtatcgtggataaccataacgggatgctgga<br>gcttggcaccagcgagcggggcgggctttccattcgcgcctggctgccagtgccggtaacgcgggcgcagggc<br>acgacaaaagaagggtaaACCTCCCAATAAGGTACCtaaGTGTcGCTGCcGAACagcN<br>NBBDDBBVVHHDDBBVVHNDDNNaggAGAAGgAGACCAGCgca |
| **Frg_C - Backbone** | CACCTCGGTCTcAAGAAGAGCgcacGACGTcaCgtCgcaGAATTCcttttcggggaaatgt<br>gcgcggaacccgggtaataataatctagaccaggcatcaaataaaacgaaaggctcagtcgaaagactggg<br>cctttcgttttatctgttgtttgtcggtgaacgctctctactagagtcacactggctcaccttcgggtgggcctttctgcgtt<br>tataggtaccaggggatcctctagagtcgacctgcaggcatgcaagcttagcaagcgaaccggaattgccagct<br>ggggcgccctctggtaaggttgggaagccctgcaaagtaaactggatggctttcttgccgccaaggatctgatgg<br>cgcaggggatcaagatctgatcaagagacaggatgaggatcgtttcgtaaattagtagcccgcctaatgagcgg<br>gcttttttttaattcccctatttgtttatttttctaaatacattcaaatatgtatccgctcatgagacaataaccctgataaat<br>gcttcaataatattgaaaaaggaagagtatgagcattcagcattttcgtgtggcgctgattccgtttttgcggcgtttt<br>gcctgccggtgtttgcgcatccggaaaccctggtgaaagtgaaagatgcggaagatcaactgggtgcgcgcgt<br>gggctatattgaactggatctgaacagcggcaaaattctggaatcttttcgtccggaagaacgttttccgatgatga<br>gcacctttaaagtgctgctgtgcggtgcggttctgagccgtgtggatgcgggccaggaacaactgggccgtcgta<br>ttcattatagccagaacgatctggtggaatatagcccggtgaccgaaaaacatctgaccgatggcatgaccgtg<br>cgtgaactgtgcagcgcggcgattaccatgagcgataacaccgcggcgaacctgctgctgacgaccattggcg<br>gtccgaaagaactgaccgcgtttctgcataacatgggcgatcatgtgacccgtctggatcgttgggaaccggaac<br>tgaacgaagcgattccgaacgatgaacgtgataccaccatgccggcagcaatggcgaccaccctgcgtaaac<br>tgctgacgggtgagctgctgaccctggcaagccgccagcaactgattgattggatggaagcggataaagtggc<br>gggtccgctgctgcgtagcgcgctgccggctggctggtttattgcggataaaagcggtgcgggcgaacgtggca<br>gccgtggcattattgcggcgctgggcccggatggtaaaccgagccgtattgtggtgatttataccaccggcagcc<br>aggcgacgatggatgaacgtaaccgtcagattgcggaaattggcgcgagcctgattaaacattggtaaaccgat<br>acaattaaaggctccttttggagccttttttttggacgaccccccagtatcagcccgtcatacttgaagctagacag<br>gcttatcttggacaagaagaagatcgcttggcctcgcgcgcagatcagttggaagaatttgtccactacgtgaaa<br>ggcgagatcaccaaggtagtaggcaaataagagctcgcttggactcctgttgatagatccagtaatgacctcag<br>aactccatctggatttgttcagaacgctcggttgccgccgggcgttttttattggtgagaatccaagcactagtaaca<br>acttatatcgtatggggctgacttcaggtgctacatttgaagagataaattgcactgaaatctagtaatattttatctga<br>ttaataagatgatcttcttgagatcgttttggtctgcgcgtaatctcttgctctgaaaacgaaaaaaccgccttgcagg<br>gcggtttttcgaaggttctctgagctaccaactctttgaaccgaggtaactggcttggaggagcgcagtcaccaaa<br>acttgtcctttcagtttagccttaaccggcgcatgacttcaagactaactcctctaaatcaattaccagtggctgctgc<br>cagtggtgcttttgcatgtctttccgggttggactcaagacgatagttaccggataaggcgcagcggtcggactga<br>acggggggttcgtgcatacagtccagcttggagcgaactgcctacccggaactgagtgtcaggcgtggaatga<br>gacaaacgcggccataacagcggaatgacaccggtaaaccgaaaggcaggaacaggagagcgcacgag<br>ggagccgccagggggaaacgcctggtatctttatagtcctgtcgggtttcgccaccactgatttgagcgtcagattt<br>cgtgatgcttgtcaggggggcggagcctatggaaaaacggctttgccgcggccctctcactttcctgttaagtatctt<br>cctggcatcttccgggaaatctccgccccgttcgtaagccatttccgctcgccgcagtcgaacgaccgagcgtag<br>cgagtcagtgagcgaggaagcggaatatatcccgaagcggcatgcatttacgttgacaccatcgaatggtgca<br>aaacctttcgcggtatggcatgatagcgcccggaagagagtcaattcagggtggtgaatgtgaaaccagtaacg |

| | |
|---|---|
| | ttatacgatgtcgcagagtatgccggtgtctcttatcagaccgtttcccgcgtggtgaaccaggccagccacgtttct<br>gcgaaaacgcgggaaaaagtggaagcggcgatggcggagctgaattacattccaaaccgcgtggcacaac<br>aactggcgggcaaacagtcgttgctgattggcgttgccacctccagtctggccctgcacgcgccgtcgcaaattgt<br>cgcggcgattaaatctcgcgccgatcaactgggtgccagcgtggtggtgtcgatggtagaacgaagcggcgtc<br>gaagactgtaaagcggcggtgcacaatcttctcgcgcaacgcgtcagtgggctgatcattaactatccgctggat<br>gaccaggatgccattgctgtggaagctgcctgcactaatgttcaggcgttatttcttgatgtctctgaccagacacca<br>atcaacagtattattttctcccatgaagacggtacgcgactgggcgtggagcatctggtcgcattgggtcaccagc<br>aaatcgcgctgttagcgggcccattaagttctgtcTCGgcgcgtctgcgtctggctggctggcataaatatctcac<br>tcgcaatcaaattcagccgatagcggaacgggaaggcgactggagtgccatgtccggttttcaacaaaccatgc<br>aaatgctgaatgagggcatcgttccaactgcgatgctggttgccaacgatcagatggcgctgggcgcaatgcgc<br>gccattaccgagtccgggctgcgcgttggtgcggatatctcggtagtgggatacgacgatacagaagacagctc<br>atgttatatcccgccgttaaccaccatcaaacaggattttcgcctgctggggcaaaccagcgtggaccgcttgctg<br>caactctctcagggccaggcggtgaagggcaatcagctgttgcccgtctcactggtgaaaagaaaaaccaccc<br>tggcgcccaatacgcaaaccgcctctccccgcgcgttggccgattcattaatgcagctggcacgacaggtttccc<br>gactggaaagcgggcagtgaggcatcaaataaaacgaaaggctcagtcgaaagactgggcctttcgttttatct<br>gttgtttgtcggtgaacgctctcctgagtaggacaaatccgccgccctagacctagggcgttcggctgcggcgag<br>cggtatcagctcactcaaaggcggtaatacgtaaatcactgcataattcgtgtagctcaaggcgcactcccgttct<br>ggataatgttttttgcgccgacatcataacggttctggctaatattctgaaatgagctgttgacaattaatcatcggctc<br>gtataatgtgtggaattgtgagcggataacaatttcacacagcactctcctaatttaaatcgcacaactcaTatgA<br>GACCAGTGCC |
